# Scaling laws in enzyme function reveal a new kind of biochemical universality

**DOI:** 10.1101/2021.02.09.430541

**Authors:** Dylan C. Gagler, Bradley Karas, Chris Kempes, Aaron D. Goldman, Hyunju Kim, Sara Imari Walker

## Abstract

All life on Earth is unified by its use of a shared set of component chemical compounds and reactions, providing a detailed model for universal biochemistry. However, this notion of universality is specific to currently observed biochemistry and does not allow quantitative predictions about examples not yet observed. Here we introduce a more generalizable concept of biochemical universality, more akin to the kind of universality discussed in physics. Using annotated genomic datasets including an ensemble of 11955 metagenomes and 1282 archaea, 11759 bacteria and 200 eukaryotic taxa, we show how four of the major enzyme functions - the oxidoreductases, transferases, hydrolases and ligases - form universality classes with common scaling behavior in their relative abundances observed across the datasets. We verify these universal scaling laws are not explained by the presence of compounds, reactions and enzyme functions shared across all known examples of life. We also demonstrate how a consensus model for the last universal common ancestor (LUCA) is consistent with predictions from these scaling laws, with the exception of ligases and transferases. Our results establish the existence of a new kind of biochemical universality, independent of the details of the component chemistry, with implications for guiding our search for missing biochemical diversity on Earth, or other for any biochemistries that might deviate from the exact chemical make-up of life as we know it, such as at the origins of life, in alien environments, or in the design of synthetic life.

## Main

Life emerges from the interplay of hundreds of chemical compounds interconverted in complex reaction networks. Some of these compounds and reactions are found across all characterized organisms ^1^, informing concepts of universal biochemistry and allowing rooting of phylogenetic relationships in the properties of a last universal common ancestor (LUCA) ^2^. Thus, universality as we have come to know it in biochemistry is a direct result of the observation that known examples of life share common details in their component compounds and reactions. However, this concept of universality arising from shared biochemistry is quite different from universality in other fields of research, such as in the physical sciences. For example, in statistical physics, universality describes properties or macroscopic features observed across large classes of systems irrespective of the specific details of any one system ^3^. Universality classes become apparent in certain limits where common patterns emerge in the ensemble statistics of large numbers of interacting component parts. In some cases, the identified universality classes can be characterized by common exponents in the power laws relating different features of a given system. When a universality class is identified with distinct exponents governing its scaling behavior, the discovery can allow predictions to guide the search for new examples, e.g. in materials discovery. Correspondingly, if biochemistry could be shown to be representative of a universality class in the physical sense, a mechanistic understanding of the identified scaling exponents could have important implications for informing models of biochemistry beyond the specific biochemistry and evolutionary history of life-as-we-know-it.

It is an open question whether or not features of biochemistry can be abstracted to demonstrate behavior consistent with characterization into a universality class (or classes), but there is good reason to suspect this might be possible. Physiology across diverse organisms is already known to follow power law scaling relationships ^4^. These are often explained by evolutionary minimization of the costs associated with hard physical limits such as those set by the laws of diffusion, gravitation, hydrodynamics, or heat dissipation ^5^. Furthermore, biochemical systems are known to display universal structure across the three phylogenetic domains in the reported scale-free (power law) connectivity of compounds *within* reaction networks ^6^. It has also been shown scaling laws apply *across* networks: the average topological properties of biochemical networks follow scaling relations across examples drawn from individuals and communities ^7^. It is therefore reasonable to conjecture that the evolution of the chemical components themselves, which compose these networks, could be subject to physical constraints that would exhibit tell-tale scaling relationships indicative of universal physical limits on their bulk properties.

Enzyme functions are a good first candidate to look at to determine if biochemistry exhibits universality in a scaling limit. Enzymes play a central role in biochemistry, catalyzing a majority of cataloged biochemical reactions. While there exist different classification systems for characterizing enzyme functions, the most thoroughly developed and widely adopted is the Enzyme Commission Classification scheme. Enzyme commission numbers organize enzyme-catalyzed reactions hierarchically, using four-digit numerical classifiers which systematically categorize enzyme function by their reaction chemistry. Each enzyme with known function is assigned an enzyme commission number for its function(s). Take for example, identifier 1.1.1.1, which is the four digit identifier for the alcohol dehydrogenase reaction. The identifier 1.x.x.x labels the class of the enzyme as oxidoreductase, 1.1.x.x specifies the subclass of oxidoreductases using CH-OH groups as electron donors, 1.1.1.x specifies the sub-subclass using CH-OH groups as electron donors with NAD+ or NADP+ as electron acceptors, and 1.1.1.1 is the specific enzyme commission number when an alcohol is the substrate (in this case the specified alcohol dehydrogenase reaction). It is important to recognize that enzyme commission numbers are assigned to enzymes to describe their function, and therefore an individual enzyme can be assigned more than one enzyme commission number depending on how many functions it has. In this way, the enzyme classification scheme provides a codified binning, or ‘coarse-graining’ in physical terms, of biochemical reaction space, where the specification of each additional digit in the enzyme commission number refers to an increasingly fine-grained specification of enzyme-catalyzed reactions.

In physics, the notion of coarse-graining is critical to identifying universality classes, because it allows ignoring most details of individual systems in favor of uncovering systematic behavior across different systems. At the coarsest scale of the first enzyme commission number digit, which corresponds to the enzyme class (EC), most details specific to individual reactions are ignored. For example in the case of oxidoreductases mentioned above, the details of the donor and acceptor do not matter: the only detail relevant to the classification of enzyme functionality as an oxidoreductase is that the reaction involves electron transfer. Biochemical reactions are grouped into seven ECs as designated by the Nomenclature Committee of the International Union of Biochemistry and Molecular Biology (NC-IUBMB) ^8^. Each of these seven classes is defined by the type of reaction catalyzed: **oxidoreductases** catalyze oxidation-reduction reactions, **transferases** catalyze the transfer of functional groups between molecules, **hydrolases** catalyze the cleavage of molecular bonds via hydrolysis, **lyases** catalyze the cleavage of bonds through means other than hydrolysis, **isomerase**s catalyze intramolecular rearrangements, **ligases** catalyze the joining of large molecules, and **translocases** catalyze the transportation of substrates across membranes. Despite this codification of biochemical reaction space and a natural interpretation in terms of the coarse-graining of catalytic function in biochemistry, there has not yet been any analyses to determine whether or not systematic trends in ECs exist across biochemical systems. However, if universality can be shown to apply across a large cross-section of biological diversity with respect to these classes, it would provide the strongest candidate yet for defining biochemical universality classes, akin to universality classes in the physical sense and allowing for the prediction of properties of unobserved or not-yet engineered biochemistries.

Herein we consider a large ensemble of biochemical systems, sampling from known biochemical diversity via the information encoded in genomes across 1282 archaea, 11759 bacteria and 200 eukaryotic taxa and 11955 metagenomes from the Department of Energy Joint Genome Institute’s Integrated Microbial Genomes and Microbiomes (DOE-JGI IMG/M) database ^9,10^. We show that common patterns emerge in the form of scaling laws at the coarse-grained, macroscopic behavior of EC diversity, with the potential to predict missing EC diversity in the biosphere. We also show how these scaling relations cannot be accounted for simply due to the presence of compounds, reactions and enzyme functions shared across known life. A consensus model of the enzyme functions in the last universal common ancestor (LUCA) ^11^ is demonstrated to be consistent with the scaling behavior of extant life for all but the ligases and transferases, suggestive that some features of the universality classes of modern biochemical systems may have emerged early on in the evolution of life on Earth. Taken as a whole, the results indicate the universality classes identified are not a direct result of the existence of component universality in the more traditional biochemical sense, and therefore is more akin to universality classes in the physical sense where the specific scaling exponents arise due to optimization against hard physical limits that would be expected to similarly constrain other examples of life in the universe, including application to synthetically designed life.

## Results

### Universal Scaling Laws Define of the Behavior of Enzyme Classes Across Diverse Biochemical Systems

We are interested in whether or not enzyme classes, as a coarse-graining of biochemical reaction space, exhibit any universal scaling behaviors, and in turn, whether or not these can reveal universality classes within subsets of biochemistry, or across all of biochemistry. We analyzed scaling patterns at the macroscopic scale of enzyme classes (EC), calculated by counting the number of unique enzyme commission identifiers with a given EC as the first digit in each dataset, see Figure 1. By focusing on enzyme classes, our analysis considers only the grouped or coarse-grained functionality of biochemical reactions, ignorant of their more detailed physiology. This coarse graining of biochemical reaction space is analogous to coarse graining in physical systems, where macroscale observables, like temperature, have been shown to be more effective (predictive) descriptions than considering the multitude of possible microstates in deriving quantitative behavior that can apply across many systems, even those not yet observed (e.g. as in how the exact kinetic motion of every particle in a gas is coarse-grained to give temperature).

**Figure 1:**
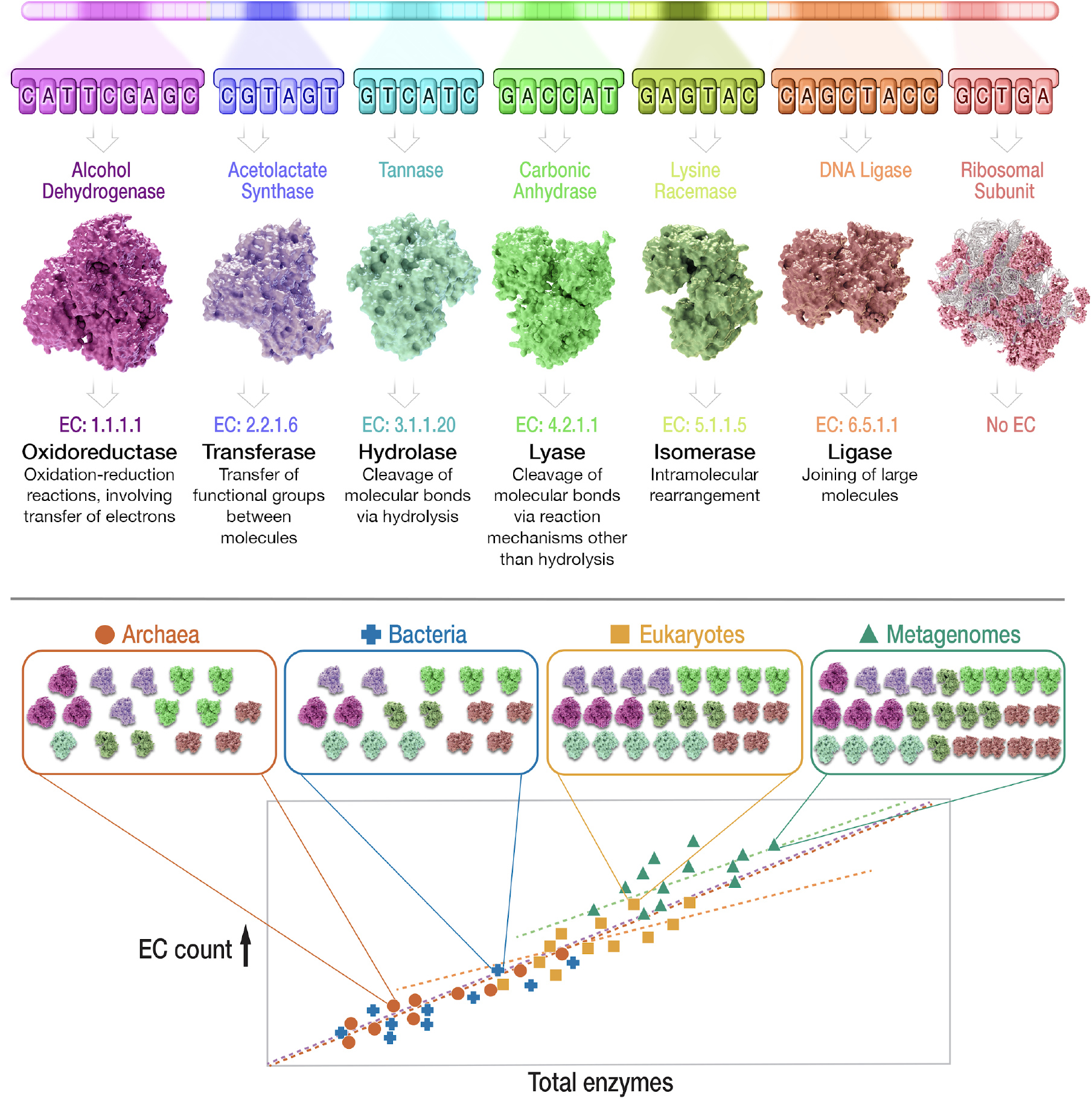
Conceptual schematic showing how genomic and metagenomic data is used to determine bulk trends in the number of enzyme functions for each major Enzyme Classes (EC). From top to bottom: In data archived by the Joint Genome Institute, many genes in genomes or metagenomes have been identified, and functions assigned to protein coding regions that are mapped to specific four digit Enzyme Commission Number identifiers. For each genomic or metagenomic sample, we then binned enzyme commission numbers based on the primary digit in the identifier, which specifies the Enzyme Class (EC): oxidoreductase (EC 1), transferase (EC 2), hydrolase (EC 3), lysase (EC 4), isomerase (EC 5), ligase (EC 6). Scaling relations are then determined by counting the total number of unique enzyme functions within a given EC (EC Count), as a function of total enzyme functions across all ECs (Total enzymes (abbreviated for Total enzyme functions)). Results are compared across the three domains archaea, bacteria and eukaryota and across metagenomes.

Each of the primary classes defined by the Nomenclature Committee of the International Union of Biochemistry and Molecular Biology (NC-IUBMB) (Webb, 1992) specifies a major class of enzymatic reactions. These include EC 1 the oxidoreductases, EC 2 the transferases, EC 3 the hydrolases, EC 4 the lysases, EC 5 the isomerases, EC 6 the ligases and EC7 the translocases (see descriptions in Figure 1). In what follows, we consider only ECs 1 through 6, as EC 7 (translocases) was only recently added, and at the time of writing is insufficiently annotated to allow performing rigorous statistical analysis across the genomes and metagenomes included in this study.

We acquired genomic and metagenomic data from the Department of Energy Joint Genome Institute’s Integrated Microbial Genomes and Microbiomes (DOE-JGI IMG/M) database ^12^. Methods for filtering the dataset to remove under- or over-annotated samples are described in the Supplement (see Section “Data Filtering”). Our final filtered data sets include 11955 metagenomes, 1282 archaea taxa, 11759 bacteria taxa and 200 eukaryotic taxa as well as 5,477 enzyme functions cataloged in the Kyoto Encyclopedia of Genes and Genomes (KEGG) database, which we use as a proxy to calculate the current estimated number of each EC in the biosphere as a whole (calculated by counting the total number of cataloged enzyme commission identifiers with that EC as the first digit in the KEGG database). By studying ensembles of biochemical systems not only at the level of individuals (genomes), but also ecosystems (metagenomes) and planetary-wide (all cataloged enzyme functions in KEGG) we are better positioned to identify universality classes in biochemistry that are scale-invariant. We are interested in scale-invariance because any scaling laws in biochemistry will be much more likely to apply to new examples of life if they are first shown to apply to known examples, independent of the scale at which we study them.

To investigate the notion that there exist size-dependent scaling relationships in EC distributions, the number of unique enzyme commission numbers in each EC class were counted for each annotated genome or metagenome in the data sets. The total number of unique enzyme commission numbers within a given EC (EC Numbers in EC Class) was then plotted as a function of the total number of unique enzyme commission numbers (Total EC Numbers) for each annotated genome or metagenome. The resultant, empirically determined scaling behaviors are shown in Figure 2, where each data point represents the binned statistics of ECs for a given genome or metagenome. These scaling relationships can be interpreted as capturing systematic changes in the number of functions within a given EC, relative to the total number of enzyme functions across all classes. We observe regular scaling behaviors for each EC across biochemical systems as they increase in size (total number of enzyme functions). Both linear regression models and power law models were fit to the data and it was determined that the power law models consistently were either equivalent to or outperformed the linear regression models using a standard error minimization test (see Supplement Section “Fitting Scaling Laws to Empirical Data”).

**Figure 2:**
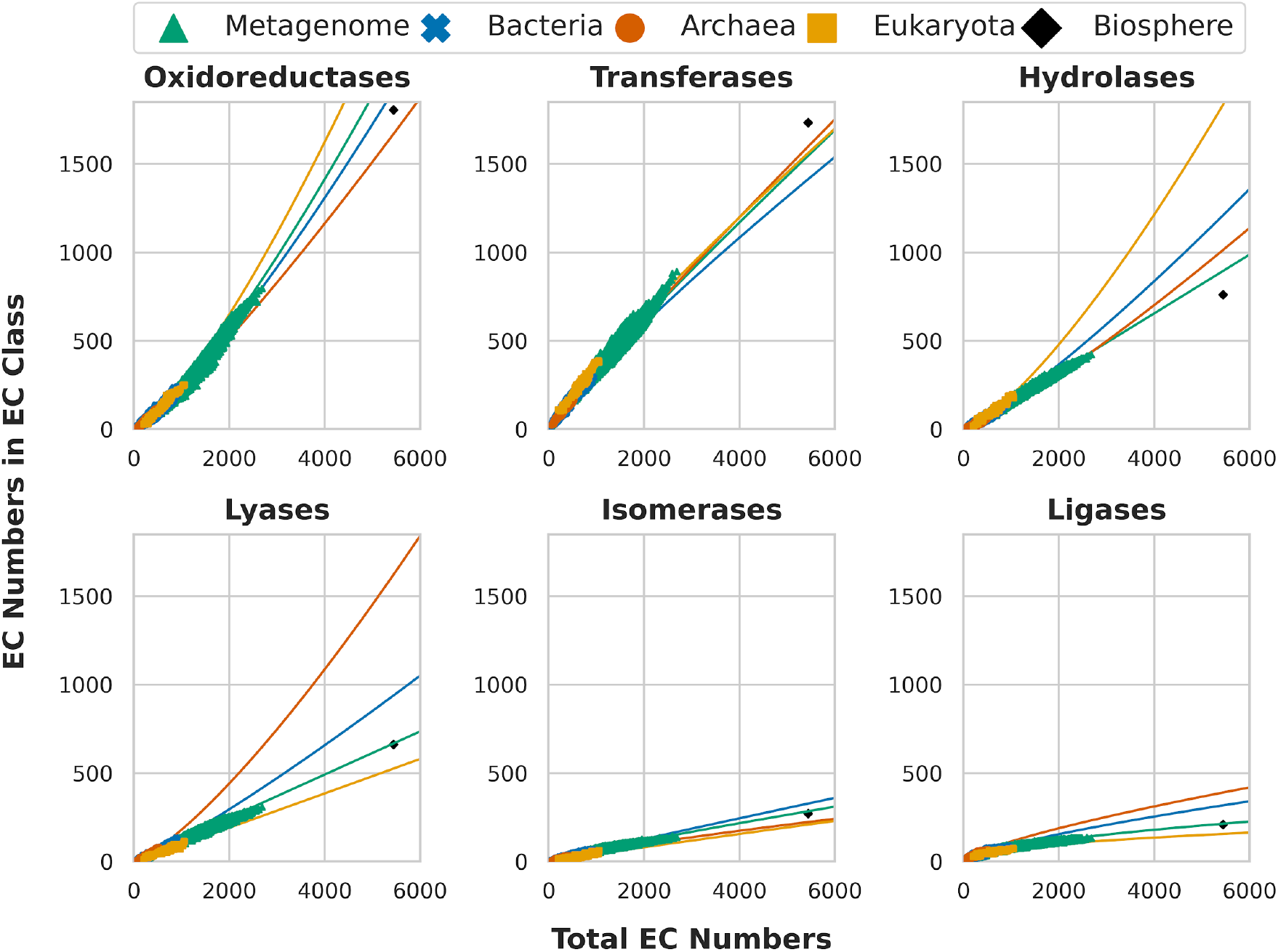
Scaling behaviors in enzyme function as captured by the number of enzyme functions within each major Enzyme Class (EC Numbers in EC Class) as a function of total enzyme functions (Total EC Numbers). Shown are the ensemble statistics for biochemical data derived from annotated genomes sampled from Archaea, Bacteria and Eukaryota taxa and from annotated metagenomes. The oxidoreductases and hydrolases display superlinear scaling across all four biochemical ensembles, whereas for transferases and ligases sublinear scaling is observed universally. The other two classes, lyases and isomerases, do not exhibit the same overall scaling trends across the different members of the ensemble (see Table 1).

We find that all ECs display scaling behavior with observed exponents greater than zero (α > 0), such that the total number of functions for each EC systematically increases with an expanding total number of enzyme functions, with some increasing much more slowly than others. Before describing these trends, it should be emphasized that many biological features do not follow scaling relationships and therefore that the observation of scaling itself is nontrivial and indicates a certain type of organizing mechanism ^5^. For example, genome size in mammals and other metazoan classes does not strongly scale with body size, and unit repair costs are roughly invariant across all of life ^13,14^. Similarly, some aspects of biochemistry that are largely conserved across all of life, such as the genetic code or translational machinery, also clearly do not exhibit scaling relationships in the diversity of their structure.

We classify the observed scaling behaviors by their scaling coefficient (with associated confidence interval) into three categories: sublinear, α < 1.0; linear, α=1.0; and superlinear, α > 1.0. Scaling behavior consistent with a linear fit (α=1.0) indicates a fixed ratio of functions within a given EC to total enzymatic functions, whereas sub- or super-linear behavior is indicative of a depletion or enrichment of functions within a given EC, respectively. Based on the sublinear, linear and superlinear classification of the empirical scaling relationships, we can determine whether specific enzyme classes display universal scaling behavior across the different data sets (three domains and metagenomes). If each dataset shares the same classification (sub-linear, linear, superlinear) for a given EC, then that grouping of functionality is a good candidate for a biochemical universality class. We are motivated to identify such classes as the strongest candidates for biochemical universality because the divide between sublinear and superlinear scaling is an important one: it can represent distinct mechanisms, or optimization for distinctly different types of constraints. ECs with the same scaling classification across all datasets therefore make possible a unified description in terms of the same underlying principles across all known examples of biochemistry. The strong case is meant to delineate those cases that make closest contact with universality classes as we know them in physics, where observations of the same properties across different systems are described by similar underlying mechanisms. Conversely, an empirically observed scaling behavior is a weak candidate for a universality class if the scaling law does not share the same classification across datasets and we do not expect these to be readily describable by universal physical limits.

Two enzyme classes, the lysases (EC 4) and isomerases (EC 5) change their classification moving from prokaryotes to eukaryotes (from superlinear to linear for lysases and from sublinear to linear in isomerases). The lysases also have different behavior for metagenomes (sublinear) meaning a strong case cannot be argued for evidence of universality classes in these ECs based on the empirically observed trends. This could be attributable to the relatively low diversity in unique functions among the lysases and isomerases as compared to other EC classes, see Table S2, meaning there may not be a sufficient number of lyases or isomerases to observe consistent trends. However, the ligases are even less diverse, yet for this class we do see universal behavior. Four classes exhibit strong evidence for universality. These include: the oxidoreductases (EC 1) and hydrolases (EC 3) both exhibiting superlinear scaling across all datasets; and transferases (EC 2) and ligases (EC 6) exhibiting sublinear scaling behavior across all data (Table 1). The results are consistent with the possibility that cataloged biochemistry is part of a universality class in each of these four enzyme classes. This introduces potential for the scaling behaviors across life to be derived from a common theory that predicts these scaling behaviors. This would be the case if, for example, all living systems are governed by the same dominant organizing constraints or principles with respect to the use of these four EC classes.

**Table 1:** Regression Values. Parameters of the logarithmic scaling law fits shown in Figure 2. Dark shaded cells indicate fits that exhibit superlinear scaling (slope > 1.0), light shaded cells indicate linear scaling (slope = 1.0), and white cells indicate sublinear scaling (slope < 1.0).

|  | Archaea | Bacteria | Eukaryota | Metagenome |
| --- | --- | --- | --- | --- |
| Oxidoreductases | 1.175+/-0.023 | 1.239+/-0.006 | 1.327+/-0.037 | 1.291+/-0.004 |
| Transferases | 0.937+/-0.013 | 0.868+/-0.003 | 0.864+/-0.021 | 0.911+/-0.003 |
| Hydrolases | 1.195+/-0.033 | 1.196+/-0.006 | 1.344+/-0.046 | 1.015+/-0.003 |
| Lyases | 1.303+/-0.022 | 1.158+/-0.005 | 1.014+/-0.046 | 0.995+/-0.003 |
| Isomerases | 0.820+/-0.028 | 0.959+/-0.006 | 0.959+/-0.066 | 0.887+/-0.004 |
| Ligases | 0.733+/-0.021 | 0.722+/-0.006 | 0.462+/-0.032 | 0.573+/-0.003 |

Within a dataset we generally observe small 95% confidence intervals on the scaling exponents (see also Tables S4-S6 for fit data, including fits to all three domains combined and all data combined which also have small confidence intervals). It should be noted that for several ECs the scaling exponents are significantly different across datasets, particularly between prokaryotes and eukaryotes. For example, the exponent for ligases, while consistently sublinear, changes by nearly 1/4 between prokaryotes and eukaryotes, which could have significant implications for differences in the diversity of polymerization reactions across different biochemical systems. The most consistent exponent values across taxa and metagenomes are for the transferases, and isomerases. We do not consider the isomerases universal because eukaryota have a coefficient more consistent with linear scaling behavior; however the error bars indicate it is also consistent with a sublinear fit for this taxa, in which case isomerases could join four of the other classes in being categorized as universal. Lysases have the largest variation in coefficients and the largest shifts in classification and therefore do not display evidence of universality in their scaling behavior. Why only the lysases really stand out in this regard is a subject of interest for future research. It could be that categorization of functions as lysases is not a natural partition of reaction space with respect to the constraints that operate on biochemical organization.

A surprising aspect of EC scaling is that the scaling relationships appear to be more universal than the previously identified scaling relationships for numerous other biological observables, which are known to change classification across domains, or across levels of organization. For example, the scaling relationships for whole-organism metabolic and growth rates are known to dramatically shift across the bacteria/eukaryote divide ^15,16^. Metabolic rates increase superlinearly with organism size for bacteria, but only linearly for unicellular eukaryotes, and sublinearly for multicellular eukaryotes ^16^. Even more strikingly, the growth rates derived from this metabolic scaling increase with cell size in bacteria but decrease with cell or body size in eukaryotes ^15^. Similarly, there are known asymptotic limits at the large end of bacteria related to increasing the number of ribosomes to keep up with growth rates, which implies another dramatic shift across this evolutionary transition ^17^. Despite these differences in other physiological observables, we find that ECs follow relatively consistent scaling behavior both across taxa and across the level of ecological organization for four of the six classes with isomerases as a possible fifth candidate. Thus, the observed EC scaling laws hold promise that missing biochemical diversity, in the form of missing enzyme functions, can be predicted from these scaling relations. For example, comparing projected trends in Figure 2 to the currently cataloged number of enzyme commission numbers in each class indicates that there are many oxidoreductase and hydrolase functions left to be discovered, whereas the scaling laws predict all ligase and isomerase functions are already known, see also Figure S6. The scaling relations underestimate biosphere-level enzyme functions only in transferase diversity, possibly due to the lower representation of eukaryota and archaea in our data set: scaling trends for these domains most closest approach the total cataloged transferase diversity.

### Universality in Scaling of Enzyme Function is not Explained by Universally Shared Components

A major challenge for any claim of universality observed across life on Earth, which seeks to inform more general principles, is the shared ancestry of known life. Evolutionary contingency influences the biochemical concept of universality because of the component set of enzyme functions, reactions and compounds common to all life, which arises due to shared evolutionary history. This allows rooting of phylogenetic relationships in the properties of a last universal common ancestor (LUCA), which is expected to share much of the same universal component set. In this sense, the biochemical concept of universality pertains to what physicists refer to as *microscale* features, which here manifests as the specific molecules and reactions used by all life. By contrast, the scaling behaviors we have identified in the previous section pertain to a *macroscale* feature arising due to the many interactions among molecules in biochemical systems. The scaling relations therefore need not, in principle, rely on the presence of a shared component chemistry across systems. In fact, in order to identify EC scaling as a potential universality class for biochemistry in the physics sense, which could allow generalizability beyond life as we know it, a key requirement is that universality in scaling behavior does not directly depend on universally-shared component chemistry. That is, it requires demonstrating the reported scaling trends are in fact genuine macroscale features and do not arise solely because of microscale properties (biochemical components) common to all life. We therefore next sought to determine the universality of component compounds, reactions, and enzyme functions across our data and in a consensus model for LUCA. Our aim is to determine whether or not the presence of universal scaling behaviors is strongly driven by the presence of universal biochemical components. We include LUCA as a model of early life, which is itself constructed based on the existence of shared components across modern systems ^11^. The consensus LUCA model we use for comparison is derived from the eight current leading models of LUCA, see Supplement Section “Consensus LUCA Enzyme Functions” for details.

Our first goal was to determine the frequency to which specific enzyme functions are distributed across our datasets. In particular, we aimed to evaluate whether the scaling trends observed in Figure 2 arise because of a set of highly redundant enzyme functions that recur across many samples within the biochemical ensemble, or are instead macroscale features that do not depend on the exact functions used across diverse biochemical systems but instead can be attributed to common patterns on how functions are used. To do so, we evaluated the component universality of each enzyme function, by rank ordering enzyme functions by their frequency of occurrence across a given data set. The results are shown in Figure 3. We then assigned Area Under the Curve (AUC) scores using Simpon’s rule to the occurrence frequencies of enzyme functions across domains and metagenomes. These AUC values allow efficient comparison of the distribution of enzyme commision numbers within a given EC for each dataset, where values closer to AUC = 1 indicate an EC has component enzyme functions that are more commonly distributed across biochemical systems and values closer to AUC = 0 indicate cases where specific enzyme functions are relatively rare across the dataset.

**Figure 3:**
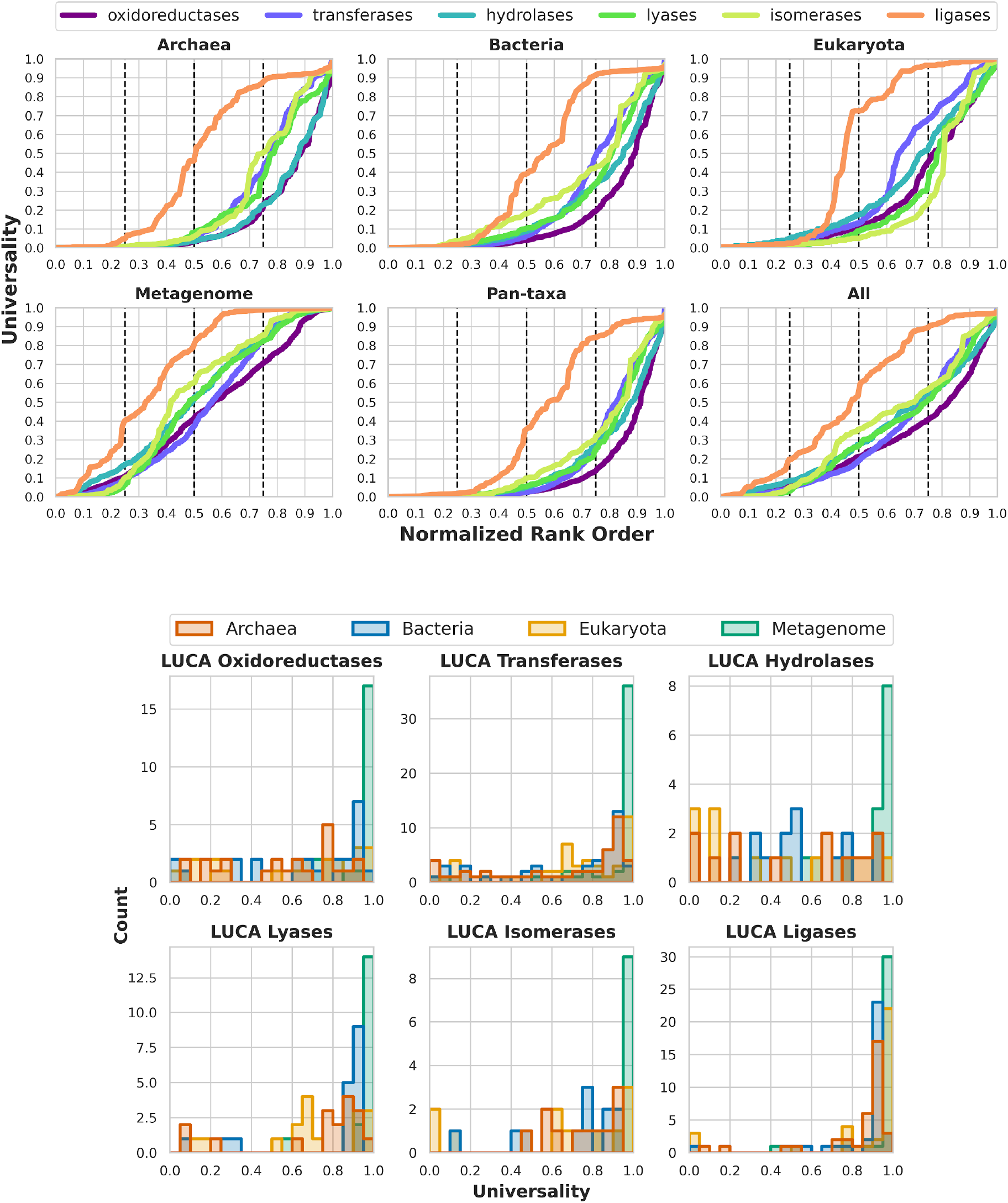
Top: Universality of specific Enzyme Commission Number identifiers for each enzyme class (EC). Specific enzyme functions are rank-ordered according to their frequency of occurrence across a given data set. Bottom: Distribution of the 154 enzyme commision numbers present in the consensus LUCA model as found across each of our datasets. The Last Universal Common Ancestor is more universally distributed across metagenomes.

AUC results for the data in Figure 3 are shown in Table 2, where “Pan-taxa” and “All” categories include data from all three domains, or from all domains and metagenomes grouped together, respectively (see also Supplement Section “Additional Scaling Plots including Pan-taxa and All Data Scaling). From the frequency of occurrence curves (Figure 3) and their AUC scores (Table 2) it is apparent that macroscale universality in the scaling behavior of ECs does not directly correlate with a high degree of microscale universality in enzyme function. This result can be seen most clearly by comparing the results in Table 2 to those in Table 1. For example, the oxidoreductases consistently exhibit the lowest AUC scores among the different enzyme classes, corresponding to a high degree of unique enzyme functions across different biochemical systems. Yet, as a class the oxidoreductases exhibit universal scaling behavior with very tightly constrained scaling coefficients, which are only second to the transferases. In contrast, the ligases have a high number of shared enzyme functions across the datasets. While the ligases are similar to oxidoreductases in that they exhibit universal scaling patterns across all data, they also have much more variation in their scaling coefficients. In fact, ligases are the most universal in terms of their components, but also have the largest variation in terms of scaling coefficient among any of the enzyme classes. Thus, the oxidoreductases and ligases give us two end-member cases. In the first, tightly constrained universal scaling behavior emerges from ensembles with relatively few universally shared component enzyme functions within the oxidoreductases. In the second, loosely constrained universal scaling behavior emerges from ensembles with a high degree of shared component ligase functions across samples. This contrast between the oxidoreductases and ligases highlights how the observed scaling trends cannot be explained directly by the universality of the underlying component enzyme commision identifiers. In fact, looking across all classes, there is no direct correlation between universality of enzyme functions within a given EC and universality in the apparent scaling behavior of the EC class as a whole (see Supplement Figure S7). This suggests that EC scaling is indeed a macroscale property that emerges because of the interactions among many individual reactions in an organism, and could reflect the presence of universal physical constraints on the architecture of biochemical networks in terms of catalytic functional diversity. It should be noted that metagenomes have the highest AUC scores, meaning it is more likely to find common enzyme functions, reactions and compounds across community samples than across individuals sampled from domains. However, we also observe metagenomes share similar EC scaling to individuals for some ECs (a key part of our interpretation of universality classes), further corroborating a lack of direct correlation between scaling patterns and AUC scores.

**Table 2:** Area Under the Curve (AUC) scores for data shown in Figure 3 A score of AUC=1.0 means an enzyme function occurs in 100% of samples and a value of 0.0 indicates a function is found in none. Thus, AUC scores closer to 1 indicate more universality in the distribution of specific functions in a given class. Shading indicates superlinear, linear or sublinear scaling behavior observed for a given class across a given data set as in Table 1.

| Domain | Oxidoreductases | Transferases | Hydrolases | Lyases | Isomerases | Ligases |
| --- | --- | --- | --- | --- | --- | --- |
| Archaea | 0.152 | 0.244 | 0.158 | 0.228 | 0.248 | 0.461 |
| Bacteria | 0.156 | 0.249 | 0.210 | 0.233 | 0.284 | 0.431 |
| Eukaryota | 0.253 | 0.337 | 0.303 | 0.233 | 0.214 | 0.522 |
| Metagenome | 0.431 | 0.451 | 0.508 | 0.479 | 0.518 | 0.663 |
| Pan-taxa | 0.129 | 0.190 | 0.173 | 0.189 | 0.214 | 0.405 |
| All | 0.270 | 0.311 | 0.330 | 0.320 | 0.357 | 0.526 |

The notion of universal biochemistry - that all life on Earth shares a specific component set of reactions and molecules - is best captured in attempts to reconstruct the last universal common ancestor (LUCA) of all life on Earth. Therefore, we also sought to understand the relationship between the ECs in a consensus model of LUCA, and the universality class of modern biochemistry as dictated by the newly identified EC scaling relations. To approach this, we first compared the distributions of LUCA enzyme functions to their universality across modern biochemical systems, see Figure 3 (bottom panel). We find that across all six ECs the majority of component enzyme functions found in the LUCA model are also found in modern metagenomes, corroborating proposals that inferred LUCA genomes represent a population of organisms rather than a single individual organism ^18^. Comparing the three domains, the patterns vary significantly by EC. For the oxidoreductases and hydrolases the enzyme functions in LUCA are nearly uniformly distributed in terms of their distribution across modern organisms: some enzyme functions are rare and some are common. The other four enzyme classes exhibit a bias toward enzyme functions from LUCA that are more universal across modern organisms. This bias is most evident in the case of the ligases, where nearly all ligase functions in the LUCA model are found in >90% of modern organisms across the three domains. This is to be expected based on how LUCA models are currently constructed: since LUCA is phylogenetically reconstructed, or at least reconstructed from consensus across extant organisms, the commonality of ligase functions across different biochemical systems means they are more likely to be represented in LUCA models than other ECs that have fewer universal enzyme functions (e.g. the oxidoreductases).

We next compared component frequencies of enzyme functions, reactions and compounds across modern life, and component frequencies in LUCA, shown in Figure 4. Associated AUC values for the distributions shown in Figure 4 (top panel) are in Table 3. The ranking of the AUC scores consistently places AUC_compounds_ > AUC_reactions_ > AUC_enzyme_ functions across all datasets: that is, independent of domain or level of organization biochemical systems tend to share more compounds in common than they do reactions, and more reactions than enzyme functions. We compared the distribution of enzyme functions, reactions and compounds found in LUCA in terms of their universality across modern life to determine the specific components that are more or less common across our datasets, and to corroborate what specific dataset(s) are more representative of the consensus LUCA, Figure 4 (bottom panel). We find that of the 154 enzymes, 337 reactions, and 438 compounds in the consensus LUCA, there is (as should be expected based on construction) a strong bias in the distribution toward inclusion of more universally distributed components. As with EC distributions, we see the majority (> 90%) of compounds, reactions and enzyme functions in LUCA are found in >90% of metagenomes, Figure 4 (bottom panel). Among the three domains, the eukaryotes share the most universal components with LUCA. We can conclude that in terms of component biochemistry, LUCA components are more universally found in metagenomes than they are in any of the three domains, consistent with hypotheses that LUCA should best be understood as an ecosystem scale property ^18–20^. We also find that the universality of LUCA enzyme functions in modern systems depends strongly on the EC.

**Figure 4:**
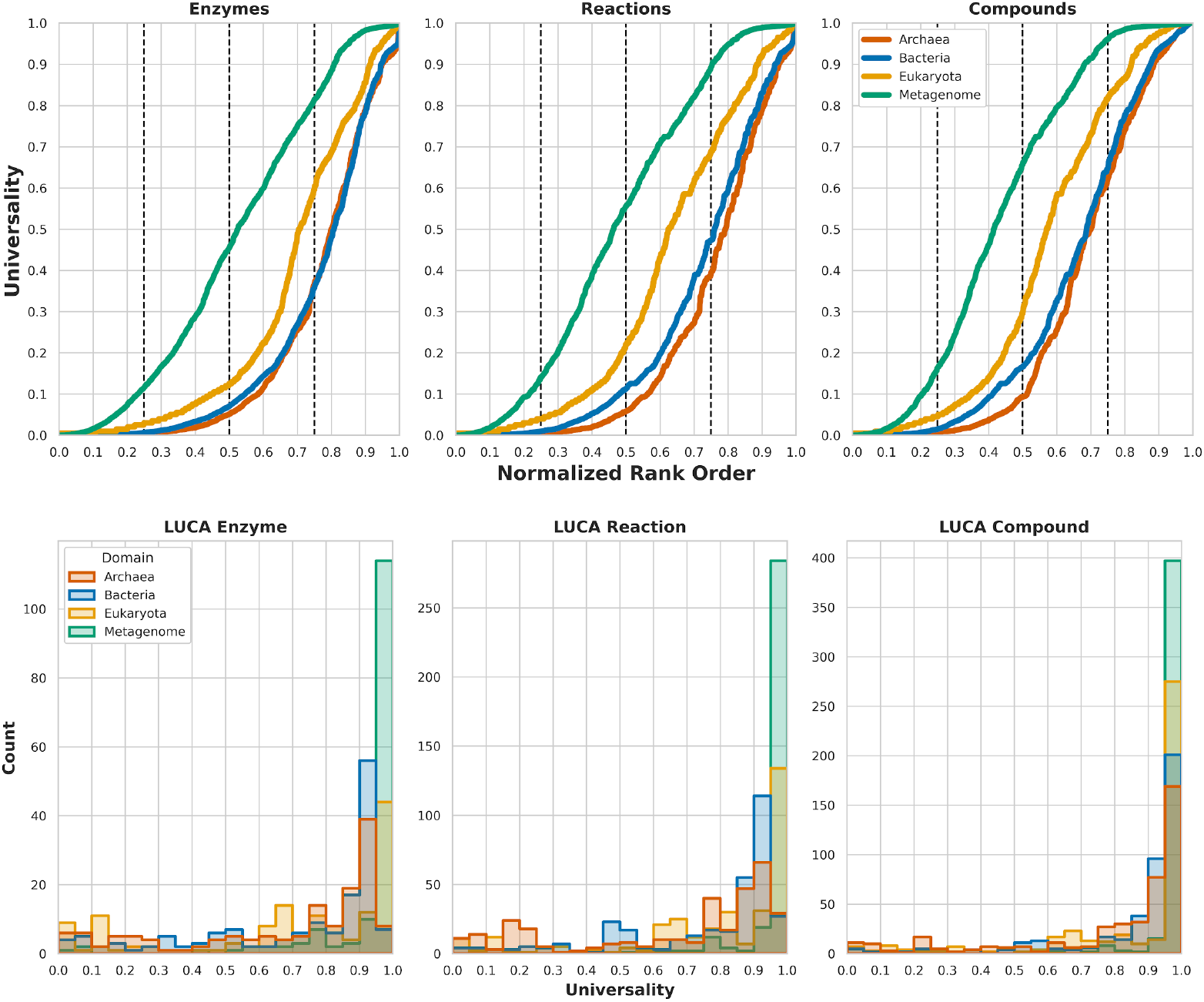
Universality curves for enzymes, reactions and compounds in modern life confirm distribution of LUCA across our datasets for modern systems.

**Table 3:** Area Under the Curve (AUC) scores for data shown in Figure 4 top panel. A score of AUC=1.0 means an enzyme function occurs in 100% of samples and a value of 0.0 indicates a function is found in none. Thus, AUC scores closer to 1 indicate more universality in the distribution of specific functions in a given category.

| Dataset | Enzyme | Reaction | Compound |
| --- | --- | --- | --- |
| Archaea | 0.220 | 0.232 | 0.307 |
| Bacteria | 0.226 | 0.266 | 0.335 |
| Eukaryota | 0.300 | 0.363 | 0.421 |
| Metagenome | 0.472 | 0.521 | 0.567 |

We also compared the EC distribution in LUCA to that of modern life. Figure 5 shows that the number of functions within each EC in LUCA falls within the scaling trends of the ensemble of modern biochemical systems, with the exception of the ligases and transferases. This could be expected for the ligases given that specific ligase functions are more universally shared across all datasets than the other ECs (see Figure 3) leading them to be heavily represented in LUCA. Our scaling fits in fact indicate the diversity of ligase functions is over-represented in LUCA relative to other classes. For the transferases, we do not see an over-representation but instead they appear under-represented based on the extrapolation of the scaling fits. In this case the interpretation is not as clear cut because the AUC scores are not consistent across datasets: AUC scores for the transferases in archaea, bacteria and metagenomes are similar in value to other EC classes, however for eukaryota the AUC score for transferases is second only to the ligases (Table 2). Nonetheless, the transferase functions present in LUCA tend to be more highly skewed toward being common in modern systems across all four datasets, much like the ligases (Figure 3, bottom).

**Figure 5:**
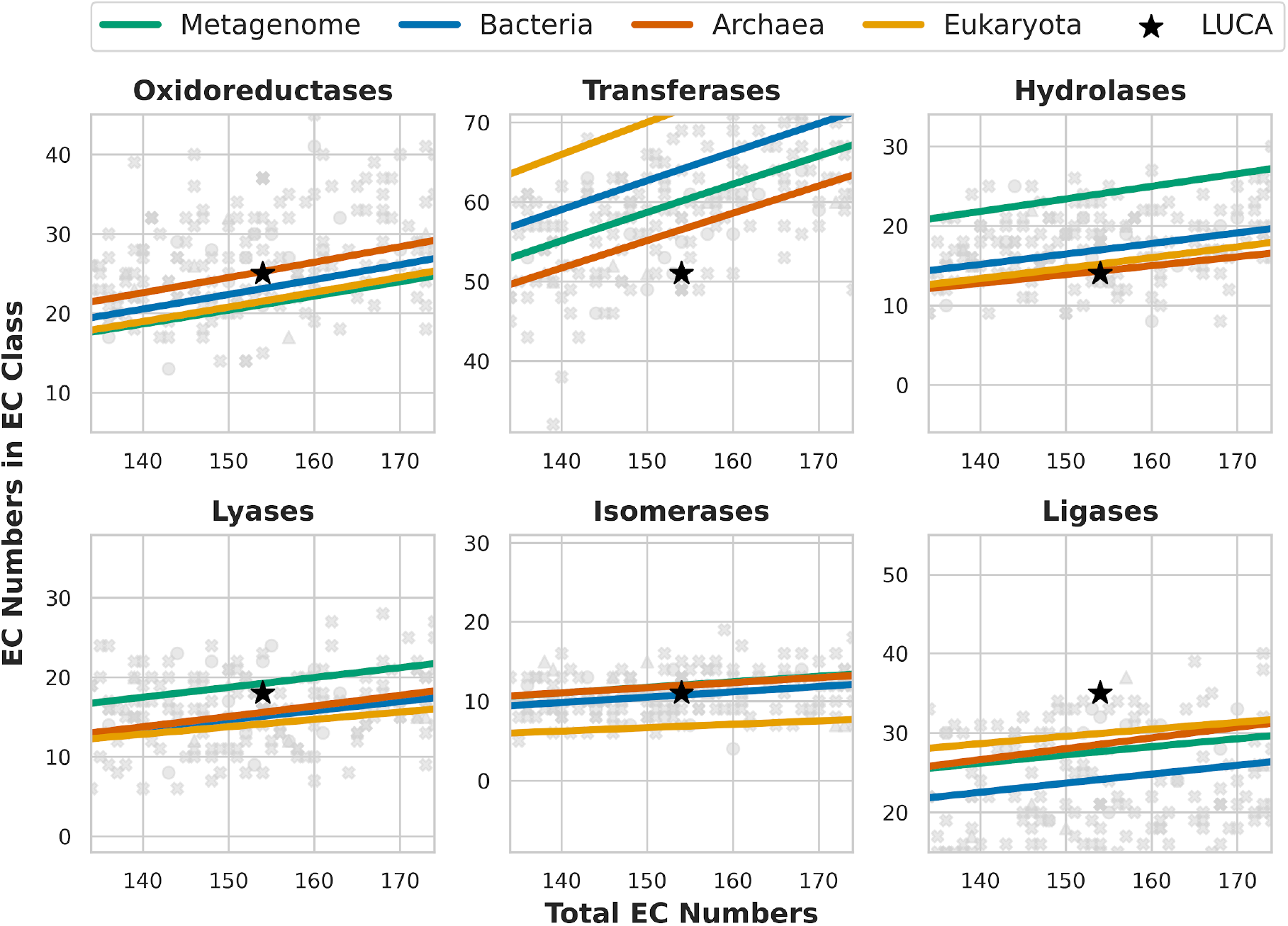
Enzyme functions in a consensus model for the Last Universal Common Ancestor (LUCA) are consistent with the universal scaling behavior of modern life for all ECs, with the exception of the ligases and transferases. For hydrolases LUCA is more consistent with an individual and for lyases an ecosystem.

For other enzyme classes, LUCA does fall within expected scaling trends. The LUCA model is most consistent with the scaling trends of individuals (domains) in terms of its diversity of hydrolases and isomerases. For isomerases this might be expected, since this EC class shows a bias toward LUCA components that are more universally distributed across modern life (Figure 3, bottom panel). For hydrolases it is more surprising: LUCA hydrolase functions are fairly randomly distributed in terms of their frequencies across modern organisms; yet the LUCA model is consistent with modern scaling trends. For lyases the situation is different, as LUCA enzyme functions in this class tend toward a ratio more consistent with the scaling trend of metagenomes. For oxidoreductases and transferases the scaling trends for both the individuals and metagenomes approach the value for LUCA. These results further corroborate that it is not a simple extrapolation from universality in component parts to observed EC scaling trends: while LUCA is most consistent with metagenomes in terms of the distribution of its component parts in modern systems, it sits on scaling curves that intersect both individuals and ecosystem ensembles or does not intersect observed scaling at all (e.g. for ligases and transferases). Surprisingly, and consistent with our hypothesis about optimization toward hard physical limits, the tightest correlation to EC scaling laws comes from those enzyme classes whose specific functions are less commonly distributed in modern biochemical systems.

## Discussion

Traditional views on the universal nature of biochemistry have focused on the existence of specific component compounds, reactions, and enzyme functions, which are conserved across known examples of life ^1^. This perspective should be thought of as universality of component membership or composition. It has informed many fields of inquiry, including efforts to constrain the chemistry implicated in the origins of life and the search for life on other worlds ^21^. This is despite the fact that we currently do not have concrete scientific inroads to understand whether the chemistry of life should or should not have the hallmark property of universality in component membership across all examples (known and unknown). However, synthetic biology is already demonstrating that perhaps component membership is not enough, given experimental evidence of alternative chemistries that can function in vitro and in vivo ^22,23^. In the current work, we have identified new systematic regularities in the form of scaling laws that allow a different window into universal biochemistry. These are more generalizable because they do not depend on the details of components and allow asking new questions about what other biochemistries could be possible, with the constraint they are consistent with predicted scaling trends.

Microscale details in the specific enzyme functions used by life can vary significantly from system to system; however, we have shown the macroscale patterns that emerge from coarse-graining enzyme functions follow tightly constrained power laws. This suggests a universal macroscale pattern in function across known life that is not dependent on the component compounds, reactions or specific enzyme functions. The universality of EC scaling is therefore likely to arise due to hard physical constraints, where the operations catalyzed among compounds in living chemistries seem to be universally constrained, independent of the specific catalyst (enzyme) identities. While enzymes fall into the category of biochemical macromolecules that are themselves part of the universal set of component membership, our focus on functions is not necessarily so restricted that it need not apply solely known biochemical catalysts (e.g. enzymes). For example, many biochemical reactions have been shown to be catalyzed by alternative polymers to those used in extant life, or by cofactors in origins of life studies ^22,24^. Since our analyses refer only to the functions of catalysis and not the catalysts themselves, they are candidates for generalizing beyond the chemistry of life-as-we-know-it.

A critical question is whether or not the universality classes identified herein are a product of the shared ancestry of life. A limitation of the traditional view of biochemical universality (focused solely on component membership) is that universality can then only be explained in terms of evolutionary contingency and shared history, which challenges our ability to generalize beyond life-as-we-know-it. Indeed, a set of closely related genomes will, by definition, share a high degree of universality in specific enzyme functions. Indeed, phylogenetic effects would be a concern here too, if we were claiming that most of biochemistry is universal in terms of shared enzyme functions as these could be attributed to oversampling highly-related genomes. Instead, we showed that a majority enzyme functions are not common across our datasets and that biological universality emerges as a macroscopic property at the level of enzyme classes, which is not directly correlated to how commonly distributed specific functions are within a class. Furthermore, enzyme class universality cannot simply be explained due to phylogenetic relatedness since the range of total enzyme functions spans two orders of magnitude, evidencing a wide coverage of genomic diversity. The maximum relatedness of two very different enzyme set sizes would occur in cases where the smaller set is a perfect subset of the larger. The very nature of the scaling relationships introduces diversity through set size differences.

The possibility of universal physical constraints on biochemical architecture is most apparent for enzyme classes that are less restricted by evolutionary contingency. That is, in cases where biological systems can innovate on function (e.g. in the oxidoreductases where there is low component universality) we see tighter constraints on the empirically determined scaling behavior across domains and levels of organization. This can be contrasted with cases where evolutionary contingency plays a more significant role (e.g. in the ligases with high component universality) where we see a larger variation in observed scaling behaviors. One possible explanation is that evolutionary constraints limit optimization toward physical limits. This is likely the explanation for the behavior of the ligases, where optimization is likely constrained by historical contingency. If the trade-off between optimization and contingency is a general feature of biochemical organization, it presents a counterintuitive approach to searching for the universal laws that could govern all biochemical systems: rather than focusing on universal components, it suggests we should instead focus efforts on cases where there is maximal diversity in component membership as it is in these cases where we might be most likely to observe optimization toward the hard physical limits that could apply to any biochemical system.

Our results have implications for understanding generalizable features of biochemistry that could apply to astrobiological searches for other examples of life, including life in alien environments or synthetically designed life, or might inform generic constraints present at the origin of life on Earth. This is because the scaling laws arise due the interactions of hundreds of chemical compounds interconverted in biochemical networks and do not depend on the specific details of those reaction networks. Thus, we can conjecture that other examples of life might be subject to the same universal constraints, which arise due to physical limitations on the architecture of complex webs of chemical reactions. Steps to validate the scaling laws identified herein as truly universal should include future work focused on uncovering the underlying mechanisms. For example, it is important to understand if the observed exponents emerge from processes that are easily generalizable and connected with physical laws, or are the result of specific and contingent evolutionary trajectories. Most of the biological scaling relationships observed thus far have been connected to the former ^5^ suggesting that there is hope that the biochemical universality that we have observed here is likewise constrained by universal physics and is therefore also itself expected to be truly universal. Building mechanistic theory to explain these scaling behaviors would also allow determining whether certain exponents are indistinguishable from one another, and would help us to assess if the mechanisms are particular to life on Earth. If the identified scaling laws are indeed universal, they can also provide new frameworks for constraining inferences about the most ancient forms of life on Earth. For example, they might inform constraints on revised models for the last universal common ancestor.

Overall, our analyses indicate that it is possible to analyze questions of biochemical universality from the perspective of statistical regularities and macroscale patterns. This opens new avenues of research into features of biochemical universality that might extend to examples not currently accessible based on notions of universal biochemistry that focus strictly on the exact identity of component compounds and molecules. Such advances will in particular become increasingly important for astrobiology as there currently exist no frameworks allowing quantitative predictions about biochemistries on other worlds

