## Supplementary Materials for "Scaling laws in enzyme function reveal a new kind of biochemical universality"

### Methods

#### Acquiring Genomic and Biochemical Data

##### Genomic and Metagenomic Data from Joint Genome Institute

Using a text-mining Python script, we retrieved metadata, genome statistics data, and the EC list for a wide array of samples from the Department of Energy Joint Genome Institute's Integrated Microbial Genomes and Microbiomes (DOE-JGI IMG/M) database. IMG/M is a comparative genomics database which contains genetic and biochemical data for various biological categories, including archaea, bacteria, eukarya, and metagenomes, among others [1]. Datasets were acquired between June 18 and June 27, 2019. Genomes and metagenomes hosted by IMG are divided into the 'JGI' and 'all' subcategories. These subcategories refer to where the sample was sequenced. For this dataset, archaea and eukarya are from the 'all' category and bacteria and metagenomes are from 'JGI'. 'all' was selected for archaea and eukarya so as to maximize the amount of representatives for these domains, whereas 'JGI' was selected for bacteria and metagenomes because sufficient representation was not a concern for these groups and we therefore elected to select for consistent annotation across datasets. As our analyses show we do not see dramatically different behavior between archaea/eukaryota and bacteria/metagenomes, and therefore can conclude that the selection of the 'all' versus 'JGI' does not in the details effect the bulk trends we report here. In total, our original dataset before filtering included 1,960 archaea samples, 16,116 bacteria samples, 677 eukarya samples, and 21,667 metagenomic samples. Archaea and bacteria samples came in the form of isolates, SAGs, and MAGs. Eukarya samples came strictly from isolates. Metagenomes came from primarily environmental samples, see Figure S1, but also include data from host-associated and engineered environments. For each sample, we pulled general study metadata (which is originally amassed in the Genome OnLine Database (GOLD) and adheres to metadata standards as defined by the Genomics Standards Consortium [2], genome statistics (which includes information such as the total number of base pairs, the total number of genes, and the number of protein coding genes, for example), and a list of enzyme commision numbers.

Through IMG's annotation pipeline, protein coding genes are assigned KEGG Orthology (KO) numbers, which are subsequently linked to enzyme commission number identifiers. Sequence data is either generated internally by the JGI, gathered from GenBank, or submitted by external researchers. Before being added to the database, unannotated sequences are put through JGI's Microbial Genome Annotation Pipeline (MGAP V.4) [3], which involves structural annotation (the identification of protein-coding genes, non-coding RNAs, regulatory RNA features and binding motifs, and CRISPR elements) and then functional annotation (function prediction of protein-coding genes). Along with alignment against different protein databases, proteins are associated with KEGG Orthology (KO) terms, from which JGI assigned enzyme commission numbers are derived.

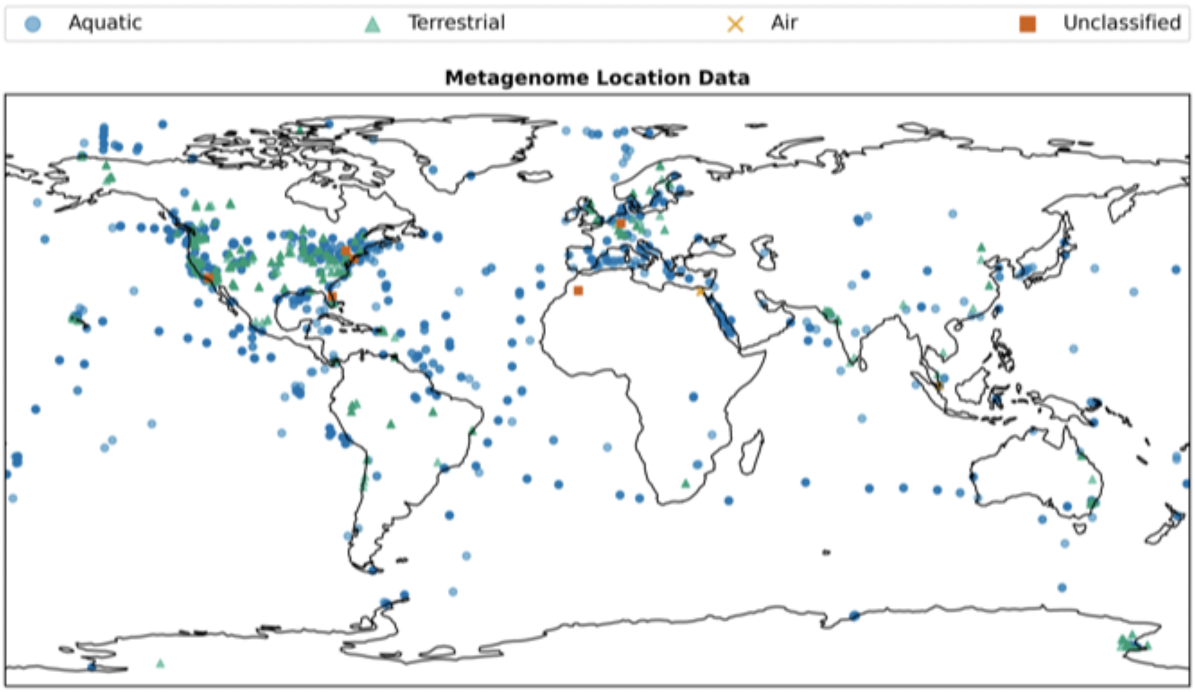


***Figure S1:*** *Distribution of the 11,955 metagenomes in our filtered dataset, including their biome classification. Data from the Joint Genome Institute [1].*

#### Biochemical Data from Kyoto Encyclopedia of Genes and Genomes

Enzyme Commission Number Identifier, reaction, and compound data were downloaded from KEGG (Kyoto Encyclopedia of Genes and Genomes) [4]. Using the enzyme commission number lists from JGI and information regarding EC/reaction and EC/compound linkages, we inferred the set of compounds and reactions that can be associated with the known enzymatic functions of each genome or metagenome.

### Coarse-graining Biochemical Space based on Enzyme Commission Number Classifications

Enzyme Commission numbers are 4-digit, hierarchical numerical classifiers which categorize enzymes based on the specific chemical reactions they catalyze [5] as designated by the Nomenclature Committee of the International Union of Biochemistry and Molecular Biology (NC-IUBMB). Enzyme commission numbers form a codified coarse-graining of biochemical reaction space, where each additional digit increases specificity about a reaction mechanism, see Table S1. Take for example, EC 1.1.1.1, alcohol dehydrogenase as highlighted in Table S1: EC 1.x.x.x refers to oxidoreductases, EC 1.1.x.x refers to oxidoreductases using CH-OH groups as electron donors, EC 1.1.1.x refers to oxidoreductases using CH-OH groups as electron donors with $NAD^+$ or $NADP^+$ as electron acceptors, and EC 1.1.1.1 refers to a specific oxidoreductase function where CH-OH groups act as the electron donors with $NAD^+$ or $NADP^+$ as electron acceptors, in this case the function is alcohol dehydrogenase. In this way, enzymes are tied to specific, experimentally verified chemical reactions. A full list of accepted enzyme commission numbers is available on the official database of the Enzyme Nomenclature List, ExplorEnz [6].

| **Hierarchical Classification Level** | **EC digit** | **Example** |
|---|---|---|
| Class | 1st digit | 1.x.x.x Oxidoreductases |
| Sub-Class | 2nd digit | 1.1.x.x CH-OH groups as donors |
| Sub-subclass | 3rd digit | 1.1.1.x NAD+ or NADP+ as electron acceptors |
| Serial Number | 4th digit | 1.1.1.1 alcohol dehydrogenase |

***Table S1:*** *Enzyme commission numbers are organized hierarchically, forming a codified coarse-graining of biochemical reaction space.*

Enzyme commission numbers are particularly appealing for this study because they classify enzymes by their reactions, as opposed to by sequence similarity or molecular structure, which offers a way to quantitatively map and categorize biochemical space from the perspective of chemical function. The primary drawbacks of the classification of enzyme functions into the hierarchy of enzyme commision numbers are that they do not include detailed information on reaction mechanisms, kinetic parameters, and small differences in sequence and protein structure [6]. However, these drawbacks do not impact our use of enzyme commission numbers in this study since we are interested in statistical patterns in the functions that this classification codifies.

Enzyme commission numbers fall into seven primary classes as designated by the Nomenclature Committee of the International Union of Biochemistry and Molecular Biology (NC-IUBMB) [5]: oxidoreductases, transferases, hydrolases, lyases, isomerases, ligases, and the newly minted translocases, see Table S2. Each of these classes is defined by the type of reaction catalyzed: oxidoreductases catalyze oxidation-reduction reactions, transferases catalyze the transfer of functional groups between molecules, hydrolases catalyze the cleavage of molecular bonds via hydrolysis, lyases catalyze the cleavage of bonds through means other than hydrolysis and typically form a double or triple bond in the process, isomerases catalyze intramolecular rearrangements, ligases catalyze the joining of large molecules, and translocases catalyze the transportation of substrates across membranes. Translocases, EC7, are a relatively new addition to the EC nomenclature and are yet to be fully integrated into mainstream biological analyses. Due to their lack of inclusion in most available data, translocases were excluded from this study.

Importantly for the current work, enzyme function can be assigned to genetic sequences via enzyme commission numbers. Once a genome or metagenome has been sequenced and protein coding regions have been identified, these protein coding regions can be referenced against protein databases to assign their function(s). Functional annotation in this study was conducted using data in the Kyoto Encyclopedia of Genes and Genomes (KEGG) [4], where proteins are assigned KEGG Orthology (KO) that contain an internal linkage in the database to enzyme commission numbers.

#### Description of Enzyme Classes

Each of the 6 primary enzyme classes used in this study have different roles in biology. As of May 2020, there are 7736 EC numbers in circulation (7646 if you exclude EC7) as per the ExplorEnz database (McDonald et al., 2009): 2402 oxidoreductases, 2085 transferases, 1783 hydrolases, 815 lyases, 326 isomerases, and 235 ligases. If you only include current enzyme commission numbers i.e. those that have not been transferred to different numbers or deleted, the numbers are as shown in Table S2, which provides the total number of enzymes in each EC class used for this study. We next briefly describe functions in each EC class.

| Class | Class Name | Function | Current Total |
|---|---|---|---|
| **1** | Oxidoreductase | Oxidation-reduction reactions, involving transfer of electrons | 1905 |
| **2** | Transferase | Transfer of functional groups between molecules | 1917 |
| **3** | Hydrolase | Cleavage of molecular bonds via hydrolysis | 1315 |
| **4** | Lyase | Cleavage of molecular bonds via reaction mechanisms other than hydrolysis | 705 |
| **5** | Isomerase | Intramolecular rearrangement | 304 |
| **6** | Ligase | Joining of large molecules | 220 |

***Table S2****: Biochemical reaction classes as defined by the primary digit of Enzyme Commission (EC) numbers as established by the Nomenclature Committee of the International Union of Biochemistry and Molecular Biology (NC-IUBMB) (EC 7 translocases, which transport substrates across membranes are not included in this study as they are a relatively new class with less annotated data then ECs 1 - 6). Totals from ExploreEnz database [6].*

##### Oxidoreductases

Oxidoreductases (EC1) are an abundant and diverse class of enzymes which catalyze oxidation-reduction reactions. Oxidoreductases are critically involved in both aerobic and anaerobic respiration as membrane electron transfer proteins, helping to drive the generation of proton gradients and subsequent ATP synthesis [7]. Furthermore, oxidoreductases are integrally involved in metabolite processing (via the synthesis of alcohols, ammonia, carboxylic acids, alkenes and alkanes, and breakdown of amino acids) and the uptake and elemental fixation of various inorganic compounds including sulfur and nitrogen [8].

##### Transferases

Transferases (EC2) are also abundant and diverse in the biosphere, catalyzing the transfer of functional groups between molecules. These common enzymes drive much of biology's core anabolic work, such as peptide bond formation, nucleotide synthesis, carbohydrate synthesis, and DNA and RNA synthesis, in addition to also participating in glycolysis, the pentose phosphate pathway, and many regulatory functions such as detoxification, molecular trafficking, protein degradation, cellular energy maintenance, and cofactor coordination [9].

##### Hydrolases

Hydrolases (EC3) catalyze the hydrolytic cleavage of various types of bonds and power much of biology's catabolic work: cleaving ester bonds (which store fatty acids as glycerides), glycosidic bonds (which compose complex sugars and are structurally involved in DNA), ether bonds (which impart biomolecules with a high degree of resistance to biological mineralization) [10], peptide bonds, and phosphate bonds [11], among others.

##### Lyases

Lyases (EC4) catalyze bond cleavage by means other than hydrolysis and differ stoichiometrically in that two or more substrates are used for one reaction direction. While hydrolases use water to cleave ester-type linkages, lyases tend to break or create double bonds, primarily carbon-carbon double bonds. Among other things, these enzymes are responsible for liberating, and thus rendering bioavailable, certain compounds like water, ammonia, chloride, and sulfide.

##### Isomerases

Isomerases (EC5) are the second least abundant enzyme class and catalyze intramolecular rearrangements. Isomerases are primarily involved in carbohydrate metabolism [12] and in terpenoid/polyketide metabolism, which is important for generating secondary metabolites [13]. Furthermore, racemases/epimerases, the primary isomerase subclass (EC 5.1.x.x), are involved in the stereochemical interconversion of amino acids, which is important for bacterial cell wall assembly and other bacterial structures and functions. Furthermore, broad spectrum racemases, which produce and release several D-amino acids to the environment, suggest the importance of isomerases in ecosystem level function [14].

##### Ligases

Ligases (EC 6) are anabolic enzymes that catalyze the joining of molecules, hydrolyzing a nucleotide triphosphate (NTP) such as ATP in the process. Ligases are involved in many biologically essential tasks, including DNA and RNA repair and regulation, coenzyme A manipulation, and protein synthesis (via aminoacyl-tRNA synthetases adding amino acids to tRNA molecules) [15].

#### Data Filtering

To make the annotation quality of the samples consistent across our dataset for the analyses in the paper, we selected a subset of the dataset available from JGI through the following steps described below.

In the enzyme commission number assignment pipeline, enzyme functions are assigned as a *subset* of protein-coding genes with function assignment, so the number of genes assigned an enzyme commission number should necessarily always be less than or equal to the number of protein function assignments. Samples that don't satisfy this condition were removed from our dataset.

Additionally, a significant portion of the metagenomic samples (~1/3) had functional assignments for 100% of their protein-coding genes. Complete functional annotations do not exist for even the best-studied organisms, such as *E. coli* and yeast. We, therefore, removed metagenomic samples with 100% functional annotation from the dataset as these are likely produced in error and are over-annotated.

Finally, we cleaned the dataset by removing samples from the dataset that fell under a threshold for their number of genes and their genome size (in base pairs). Thresholds were determined by searching for information about gene counts and genome size in minimal genomes/organisms. For archaea and bacteria, we removed samples with fewer than 1,364 genes, which is reflective of sizes of the smallest free-living prokaryotes, from which we selected Pelagibacter ubique with 1,354 genes [16] as a representative size. For eukarya, we removed samples with fewer than 4,718 genes, based on the size of one of the smallest known free-living eukaryotes, Ashbya gossypii [17].

Unlike genome datasets, metagenomes were a bit more complicated for the filtering since there is no coherent concept of genome length for metagenomes or of a "minimal" metagenome. We removed metagenomes with fewer than 20,000 genes, leaving space for the smallest metagenomes to hypothetically contain some 10-20 individual archaea or bacteria.

The final product was a dataset containing 1,194 archaea (a 36.5% reduction), 10,434 bacteria (a 29.4% reduction), 267 eukarya (a 60.5% reduction), and 6,112 metagenomes (a 44.6% reduction). See Table S3 for the statistics on the initial and cleaned datasets and Figures S2 - S5 for statistical distributions of raw and filtered data.

| | **Domain** | **Initial** | **No enzyme** | **Filtered** |
|---|---|---|---|---|
| Raw | Archaea | 1960 | 0 | 1960 |

| | | | | |
|---|---|---|---|---|
| Filtered | **Archaea** | 1960 | 0 | 1282 |
| Raw | Bacteria | 16755 | 639 | 16116 |
| Filtered | **Bacteria** | 16755 | 150 | 11759 |
| Raw | Eukaryota | 710 | 33 | 677 |
| Filtered | **Eukaryota** | 710 | 0 | 200 |
| Raw | Metagenome | 21667 | 0 | 21667 |
| Filtered | **Metagenome** | 21667 | 0 | 11955 |

***Table S3:*** *Statistics for raw versus filtered data sampled from JGI.*

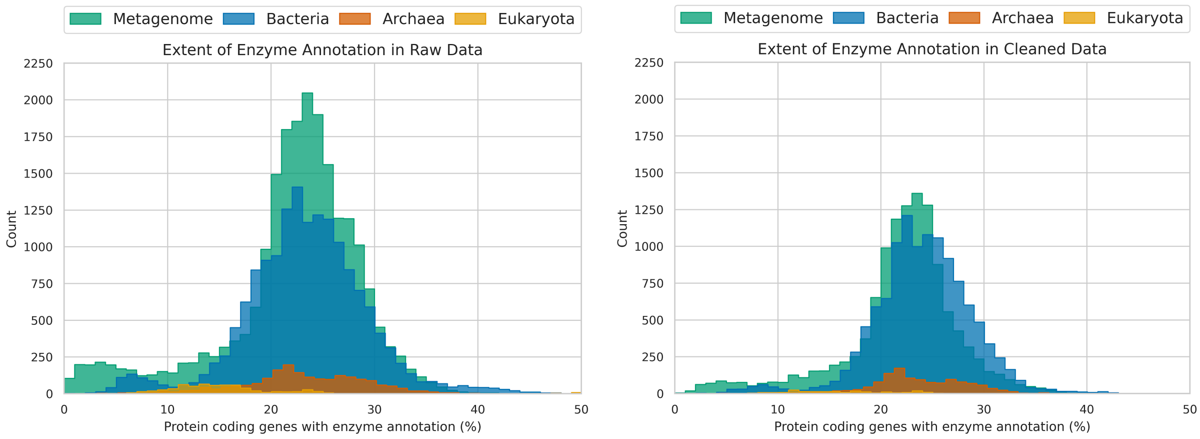


***Figure S2:*** *Extent of enzyme annotation in raw (left) and filtered (right) data sets. Histogram plots with bins of 1% width showing the extent of enzyme annotation in our dataset prior to any data cleaning or processing on the left and after filtering on the right. The x-axis depicts the percentage of protein coding genes with enzymatic assignments out of total protein coding genes in a given genome or metagenome. Enzyme assignments are made via KEGG Orthology (KO).*

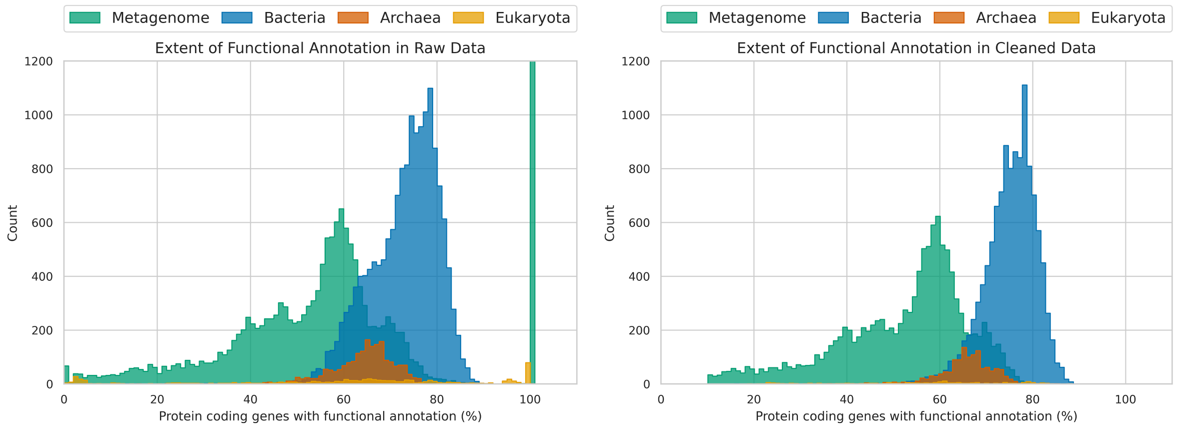


***Figure S3:*** *Extent of functional annotation in raw (left) and filtered (right) datasets. Histogram plots with bins of 1% width showing the extent of functional annotation in our dataset prior to any data cleaning or processing on the left and after filtering on the right. The x-axis depicts the percentage of protein coding genes with functional assignments out of total protein coding genes in a given genome or metagenome. Note the substantial fraction of metagenomes with ~100% functional annotation (N = 7591 with an additional N = 11 of > 120%), which are not viable for use due to artifacts of over-annotation.*

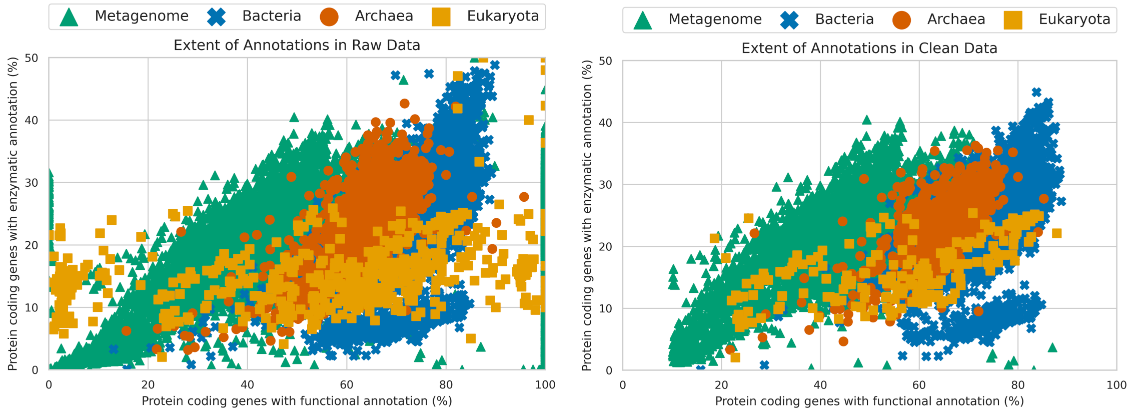


***Figure S4:*** *Extent of protein coding genes with enzymatic versus functional annotation in raw (left) and filtered (right) datasets. Scatter plot showing the extent of annotation in the dataset prior to any data processing or filtering on the left and after filtering on the right. The x-axis depicts the percentage of protein coding genes with functional annotation in each genome or metagenome and the y-axis depicts the percentage of protein coding genes with enzymatic annotation in each genome or metagenome. Note the eukarya data with enzyme annotation percentages between 0 and 25% and functional annotation percentages less than 20%. Fungi were removed from the dataset (see description in text).*

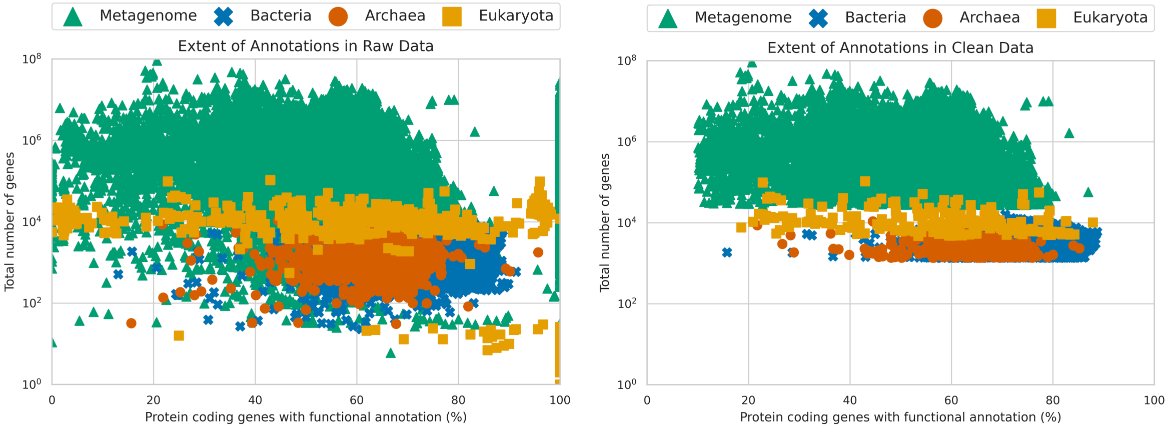


***Figure S5:*** *Extent of protein coding genes with enzymatic versus functional annotation in raw (left) and filtered (right) datasets. Scatter plot showing the extent of annotation in the dataset prior to any data processing or cleaning on the left and after filtering on the right. The x-axis depicts the percentage of protein coding genes with functional annotation in each genome or metagenome and the y-axis depicts the total number of genes in the genome. Enzyme annotations are assigned via functional annotations, so y-axis values that are greater than their corresponding x-axis values are not sensible. Fungi were removed from the dataset also.*

### Consensus LUCA Enzyme Functions

In order to identify consensus enzyme function predictions for the last universal common ancestor (LUCA), the results of eight previously published LUCA genome studies [18–25] were mapped onto clusters within the Eggnog database [26]. LUCA genome predictions from all eight studies were mapped onto UniProt accession [27] as in the LUCApedia database [28]. These UniProt accessions represent individual proteins, but the genome content of LUCA is more appropriately represented in modern taxa as larger protein families. As such, the UniProt accessions corresponding to the results of each LUCA genome study were mapped onto protein families in the EggNOG database by way of the file, <uniprot-15-May-2015.LUCA.tsv>, downloaded from the EggNOG FTP site. Any protein family predicted by four or more of the eight LUCA genome studies was retained as a consensus LUCA protein family, which resulted in 366 such families.

An ancestral enzyme commission number was subsequently inferred for each consensus LUCA protein family. First, the enzyme commission numbers associated with each consensus LUCA protein family were identified through the annotations of their component proteins found in the Uniprot database. Only reviewed Uniprot accessions were considered in this analysis [29]. Of the 366 consensus LUCA protein families, 310 contained at least one reviewed UniProt accession with an associated enzyme commission number. In order to infer whether a given enzyme function was ancestral to the protein family, the associated NCBI Taxonomic IDs of each UniProt accession in an EggNOG family were used to determine how common the enzyme function was across the three domains of life. If an enzyme function was only predicted in one taxonomic domain, it was not included in the final list of consensus LUCA enzyme commission numbers. If a single EggNOG cluster contained more than one associated enzyme commission number, only the enzyme commission number(s) with the broadest taxonomic range were retained. The resulting list of consensus LUCA enzyme functions contains 200 EC numbers from 199 Eggnog clusters. The attached appendix contains the consensus LUCA Eggnog clusters and their predicted ancestral enzyme commission number(s).

#### Fitting Scaling Laws to Empirical Data

Power laws are the natural way to address features that have consistent relationships over large changes in scale. This is in contrast to the linear or polynomial fits often used in molecular biology to capture the interconnection of features. It should be noted that a power-law fit reduces to a linear fit when the exponent is equal to one. Below we provide ordinary least squares fits for both the linear and power-law fits to the data. Our analyses show the power laws are a consistently better fit to the data and typically have exponents that are distinguishable from one, thus motivating our discussion of power law fits in the main paper.

| | **Dataset** | **Archaea** | **Bacteria** | **Eukaryota** | **Metagenome** | **Pan-taxa** | **All** |
|---|---|---|---|---|---|---|---|
| EC 1 | Slope | 0.227 | 0.266 | 0.277 | 0.345 | 0.263 | 0.312 |
| Oxidoreductases | Slope CI | -0.005 | -0.001 | -0.007 | -0.001 | -0.001 | -0.001 |
| | Intercept | -12.373 | -31.474 | -31.754 | -110.285 | -28.980 | -58.987 |
| | Intercept CI | -1.712 | -0.764 | -4.823 | -1.587 | -0.666 | -0.559 |
| | R-squared | 0.876 | 0.929 | 0.966 | 0.970 | 0.933 | 0.983 |
| | Slope Std Error | 0.002 | 0.001 | 0.004 | 0.001 | 0.001 | 0.000 |
| | Intercept Std Error | 0.873 | 0.390 | 2.446 | 0.810 | 0.340 | 0.285 |
| EC 2 | Slope | 0.311 | 0.292 | 0.338 | 0.294 | 0.301 | 0.299 |
| Transferases | Slope CI | -0.006 | -0.001 | -0.010 | -0.001 | -0.001 | 0.000 |
| | Intercept | 13.256 | 31.949 | 27.889 | 36.558 | 25.992 | 28.123 |
| | Intercept CI | -2.057 | -0.768 | -6.616 | -1.512 | -0.717 | -0.467 |
| | R-squared | 0.901 | 0.939 | 0.957 | 0.963 | 0.941 | 0.987 |

| | | | | | | | |
|---|---|---|---|---|---|---|---|
| | Slope Std Error | 0.003 | 0.001 | 0.005 | 0.001 | 0.001 | 0.000 |
| | Intercept Std Error | 1.049 | 0.392 | 3.355 | 0.771 | 0.366 | 0.238 |
| EC 3 | Slope | 0.142 | 0.171 | 0.201 | 0.156 | 0.175 | 0.168 |
| Hydrolases | Slope CI | -0.004 | -0.001 | -0.007 | 0.000 | -0.001 | 0.000 |
| | Intercept | -10.704 | -14.566 | -23.074 | 6.497 | -17.245 | -11.903 |
| | Intercept CI | -1.566 | -0.465 | -4.518 | -0.572 | -0.433 | -0.228 |
| | R-squared | 0.767 | 0.935 | 0.944 | 0.981 | 0.936 | 0.990 |
| | Slope Std Error | 0.002 | 0.000 | 0.003 | 0.000 | 0.000 | 0.000 |
| | Intercept Std Error | 0.798 | 0.237 | 2.291 | 0.292 | 0.221 | 0.116 |
| EC 4 | Slope | 0.163 | 0.136 | 0.096 | 0.118 | 0.131 | 0.123 |
| Lyases | Slope CI | -0.003 | -0.001 | -0.005 | 0.000 | -0.001 | 0.000 |
| | Intercept | -10.991 | -8.147 | -1.131 | 7.611 | -5.040 | -0.183 |
| | Intercept CI | -1.018 | -0.327 | -3.413 | -0.596 | -0.339 | -0.198 |
| | R-squared | 0.912 | 0.949 | 0.871 | 0.965 | 0.931 | 0.986 |
| | Slope Std Error | 0.001 | 0.000 | 0.003 | 0.000 | 0.000 | 0.000 |
| | Intercept Std Error | 0.519 | 0.167 | 1.731 | 0.304 | 0.173 | 0.101 |
| EC 5 | Slope | 0.050 | 0.064 | 0.045 | 0.050 | 0.063 | 0.055 |
| Isomerases | Slope CI | -0.002 | 0.000 | -0.004 | 0.000 | 0.000 | 0.000 |
| | Intercept | 6.102 | 1.197 | -1.716 | 15.035 | 1.647 | 6.407 |
| | Intercept CI | -0.737 | -0.255 | -2.507 | -0.361 | -0.248 | -0.135 |
| | R-squared | 0.645 | 0.872 | 0.734 | 0.930 | 0.854 | 0.968 |
| | Slope Std Error | 0.001 | 0.000 | 0.002 | 0.000 | 0.000 | 0.000 |
| | Intercept Std Error | 0.376 | 0.130 | 1.271 | 0.184 | 0.127 | 0.069 |
| EC 6 | Slope | 0.107 | 0.071 | 0.043 | 0.037 | 0.067 | 0.043 |
| Ligases | Slope CI | -0.003 | -0.001 | -0.003 | 0.000 | -0.001 | 0.000 |
| | Intercept | 14.711 | 21.084 | 29.786 | 44.602 | 23.657 | 36.555 |
| | Intercept CI | -1.129 | -0.348 | -2.041 | -0.318 | -0.333 | -0.161 |

| | | | | | | | |
|---|---|---|---|---|---|---|---|
| | R-squared | 0.781 | 0.816 | 0.789 | 0.905 | 0.784 | 0.928 |
| | Slope Std Error | 0.002 | 0.000 | 0.002 | 0.000 | 0.000 | 0.000 |
| | Intercept Std Error | 0.576 | 0.178 | 1.035 | 0.162 | 0.170 | 0.082 |

***Table S4:*** *Table of regression values for linear fits to scaling behavior of ECs across domains and metagenomes.*

| | Dataset | Archaea | Bacteria | Eukaryota | Metagenome | Pan-taxa | All |
|---|---|---|---|---|---|---|---|
| EC 1 | Slope | 1.175 | 1.239 | 1.327 | 1.291 | 1.229 | 1.255 |
| Oxidoreductases | Slope CI | -0.023 | -0.006 | -0.037 | -0.004 | -0.006 | -0.002 |
| | Intercept | -1.169 | -1.347 | -1.570 | -1.499 | -1.317 | -1.389 |
| | Intercept CI | -0.059 | -0.017 | -0.102 | -0.014 | -0.016 | -0.007 |
| | R-squared | 0.885 | 0.925 | 0.962 | 0.965 | 0.929 | 0.979 |
| | Slope Std Error | 0.012 | 0.003 | 0.019 | 0.002 | 0.003 | 0.001 |
| | Intercept Std Error | 0.030 | 0.009 | 0.052 | 0.007 | 0.008 | 0.003 |
| EC 2 | Slope | 0.937 | 0.868 | 0.864 | 0.911 | 0.890 | 0.901 |
| Transferases | Slope CI | -0.013 | -0.003 | -0.021 | -0.003 | -0.003 | -0.001 |
| | Intercept | -0.298 | -0.091 | -0.036 | -0.214 | -0.153 | -0.183 |
| | Intercept CI | -0.034 | -0.009 | -0.059 | -0.009 | -0.009 | -0.004 |
| | R-squared | 0.936 | 0.956 | 0.970 | 0.969 | 0.956 | 0.986 |
| | Slope Std Error | 0.007 | 0.002 | 0.011 | 0.001 | 0.002 | 0.001 |
| | Intercept Std Error | 0.017 | 0.005 | 0.030 | 0.005 | 0.004 | 0.002 |
| EC 3 | Slope | 1.195 | 1.196 | 1.344 | 1.015 | 1.241 | 1.158 |
| Hydrolases | Slope CI | -0.033 | -0.006 | -0.046 | -0.003 | -0.006 | -0.002 |
| | Intercept | -1.458 | -1.387 | -1.759 | -0.839 | -1.514 | -1.289 |
| | Intercept CI | -0.084 | -0.015 | -0.127 | -0.008 | -0.016 | -0.006 |

| | | | | | | | |
|---|---|---|---|---|---|---|---|
| | R-squared | 0.795 | 0.937 | 0.944 | 0.979 | 0.929 | 0.977 |
| | Slope Std Error | 0.017 | 0.003 | 0.023 | 0.001 | 0.003 | 0.001 |
| | Intercept Std Error | 0.043 | 0.008 | 0.065 | 0.004 | 0.008 | 0.003 |
| EC 4 | Slope | 1.303 | 1.158 | 1.014 | 0.995 | 1.124 | 1.047 |
| Lyases | Slope CI | -0.022 | -0.005 | -0.046 | -0.003 | -0.005 | -0.002 |
| | Intercept | -1.657 | -1.355 | -1.068 | -0.892 | -1.258 | -1.053 |
| | Intercept CI | -0.057 | -0.014 | -0.127 | -0.010 | -0.014 | -0.006 |
| | R-squared | 0.911 | 0.944 | 0.906 | 0.972 | 0.929 | 0.977 |
| | Slope Std Error | 0.011 | 0.003 | 0.023 | 0.002 | 0.003 | 0.001 |
| | Intercept Std Error | 0.029 | 0.007 | 0.065 | 0.005 | 0.007 | 0.003 |
| EC 5 | Slope | 0.820 | 0.958 | 0.959 | 0.887 | 0.944 | 0.921 |
| Isomerases | Slope CI | -0.028 | -0.006 | -0.066 | -0.004 | -0.006 | -0.002 |
| | Intercept | -0.718 | -1.064 | -1.263 | -0.861 | -1.029 | -0.967 |
| | Intercept CI | -0.072 | -0.017 | -0.185 | -0.013 | -0.017 | -0.007 |
| | R-squared | 0.714 | 0.885 | 0.803 | 0.938 | 0.867 | 0.959 |
| | Slope Std Error | 0.015 | 0.003 | 0.034 | 0.002 | 0.003 | 0.001 |
| | Intercept Std Error | 0.037 | 0.009 | 0.094 | 0.007 | 0.009 | 0.004 |
| EC 6 | Slope | 0.733 | 0.722 | 0.462 | 0.573 | 0.675 | 0.559 |
| Ligases | Slope CI | -0.021 | -0.006 | -0.032 | -0.003 | -0.006 | -0.002 |
| | Intercept | -0.147 | -0.197 | 0.466 | 0.189 | -0.063 | 0.242 |
| | Intercept CI | -0.052 | -0.016 | -0.088 | -0.010 | -0.016 | -0.006 |
| | R-squared | 0.794 | 0.834 | 0.807 | 0.914 | 0.801 | 0.907 |
| | Slope Std Error | 0.010 | 0.003 | 0.016 | 0.002 | 0.003 | 0.001 |
| | Intercept Std Error | 0.027 | 0.008 | 0.045 | 0.005 | 0.008 | 0.003 |

***Table S5:*** *Table of regression values for power law (log base 10) fits to scaling behavior of ECs across domains and metagenomes.*

| | **Dataset** | **Archaea** | **Bacteria** | **Eukaryota** | **Metagenome** | **Pan-taxa** | **All** |
|---|---|---|---|---|---|---|---|
| EC 1 | Slope | 1.175 | 1.239 | 1.327 | 1.291 | 1.229 | 1.255 |
| Oxidoreductases | Slope CI | -0.023 | -0.006 | -0.037 | -0.004 | -0.006 | -0.002 |
| | Intercept | -2.692 | -3.101 | -3.616 | -3.452 | -3.032 | -3.197 |
| | Intercept CI | -0.136 | -0.040 | -0.236 | -0.032 | -0.036 | -0.015 |
| | R-squared | 0.885 | 0.925 | 0.962 | 0.965 | 0.929 | 0.979 |
| | Slope Std Error | 0.012 | 0.003 | 0.019 | 0.002 | 0.003 | 0.001 |
| | Intercept Std Error | 0.069 | 0.020 | 0.120 | 0.016 | 0.018 | 0.008 |
| EC 2 | Slope | 0.937 | 0.868 | 0.864 | 0.911 | 0.890 | 0.901 |
| Transferases | Slope CI | -0.013 | -0.003 | -0.021 | -0.003 | -0.003 | -0.001 |
| | Intercept | -0.687 | -0.209 | -0.082 | -0.492 | -0.353 | -0.421 |
| | Intercept CI | -0.079 | -0.021 | -0.135 | -0.021 | -0.020 | -0.009 |
| | R-squared | 0.936 | 0.956 | 0.970 | 0.969 | 0.956 | 0.986 |
| | Slope Std Error | 0.007 | 0.002 | 0.011 | 0.001 | 0.002 | 0.001 |
| | Intercept Std Error | 0.040 | 0.011 | 0.069 | 0.011 | 0.010 | 0.005 |
| EC 3 | Slope | 1.195 | 1.196 | 1.344 | 1.015 | 1.241 | 1.158 |
| Hydrolases | Slope CI | -0.033 | -0.006 | -0.046 | -0.003 | -0.006 | -0.002 |
| | Intercept | -3.357 | -3.194 | -4.051 | -1.931 | -3.487 | -2.968 |
| | Intercept CI | -0.194 | -0.035 | -0.293 | -0.019 | -0.036 | -0.015 |
| | R-squared | 0.795 | 0.937 | 0.944 | 0.979 | 0.929 | 0.977 |
| | Slope Std Error | 0.017 | 0.003 | 0.023 | 0.001 | 0.003 | 0.001 |
| | Intercept Std Error | 0.099 | 0.018 | 0.149 | 0.010 | 0.018 | 0.008 |
| EC 4 | Slope | 1.303 | 1.158 | 1.014 | 0.995 | 1.124 | 1.047 |
| Lyases | Slope CI | -0.022 | -0.005 | -0.046 | -0.003 | -0.005 | -0.002 |

| | | | | | | | |
|---|---|---|---|---|---|---|---|
| | Intercept | -3.816 | -3.119 | -2.459 | -2.055 | -2.896 | -2.425 |
| | Intercept CI | -0.130 | -0.032 | -0.293 | -0.022 | -0.033 | -0.013 |
| | R-squared | 0.911 | 0.944 | 0.906 | 0.972 | 0.929 | 0.977 |
| | Slope Std Error | 0.011 | 0.003 | 0.023 | 0.002 | 0.003 | 0.001 |
| | Intercept Std Error | 0.066 | 0.016 | 0.149 | 0.011 | 0.017 | 0.007 |
| EC 5 | Slope | 0.820 | 0.958 | 0.959 | 0.887 | 0.944 | 0.921 |
| Isomerases | Slope CI | -0.028 | -0.006 | -0.066 | -0.004 | -0.006 | -0.002 |
| | Intercept | -1.653 | -2.450 | -2.909 | -1.982 | -2.369 | -2.228 |
| | Intercept CI | -0.166 | -0.039 | -0.426 | -0.030 | -0.039 | -0.016 |
| | R-squared | 0.714 | 0.885 | 0.803 | 0.938 | 0.867 | 0.959 |
| | Slope Std Error | 0.015 | 0.003 | 0.034 | 0.002 | 0.003 | 0.001 |
| | Intercept Std Error | 0.085 | 0.020 | 0.216 | 0.015 | 0.020 | 0.008 |
| EC 6 | Slope | 0.733 | 0.722 | 0.462 | 0.573 | 0.675 | 0.559 |
| Ligases | Slope CI | -0.021 | -0.006 | -0.032 | -0.003 | -0.006 | -0.002 |
| | Intercept | -0.340 | -0.454 | 1.073 | 0.435 | -0.144 | 0.558 |
| | Intercept CI | -0.120 | -0.036 | -0.202 | -0.023 | -0.036 | -0.015 |
| | R-squared | 0.794 | 0.834 | 0.807 | 0.914 | 0.801 | 0.907 |
| | Slope Std Error | 0.010 | 0.003 | 0.016 | 0.002 | 0.003 | 0.001 |
| | Intercept Std Error | 0.061 | 0.019 | 0.103 | 0.012 | 0.018 | 0.008 |

***Table S6:*** *Table of regression values for power law (log base e) fits to scaling behavior of ECs across domains and metagenomes.*

#### Additional Scaling Plots including Pan-taxa and All Data Scaling

In the main manuscript we report scaling laws for the individual domains and metagenomes. Also of interest is the scaling behavior of all individuals (combining data from all three domains) or the scaling behavior of all biochemical systems in our ensemble (combining data from all three domains and metagenomes). These scaling trends are shown in Figure S6.

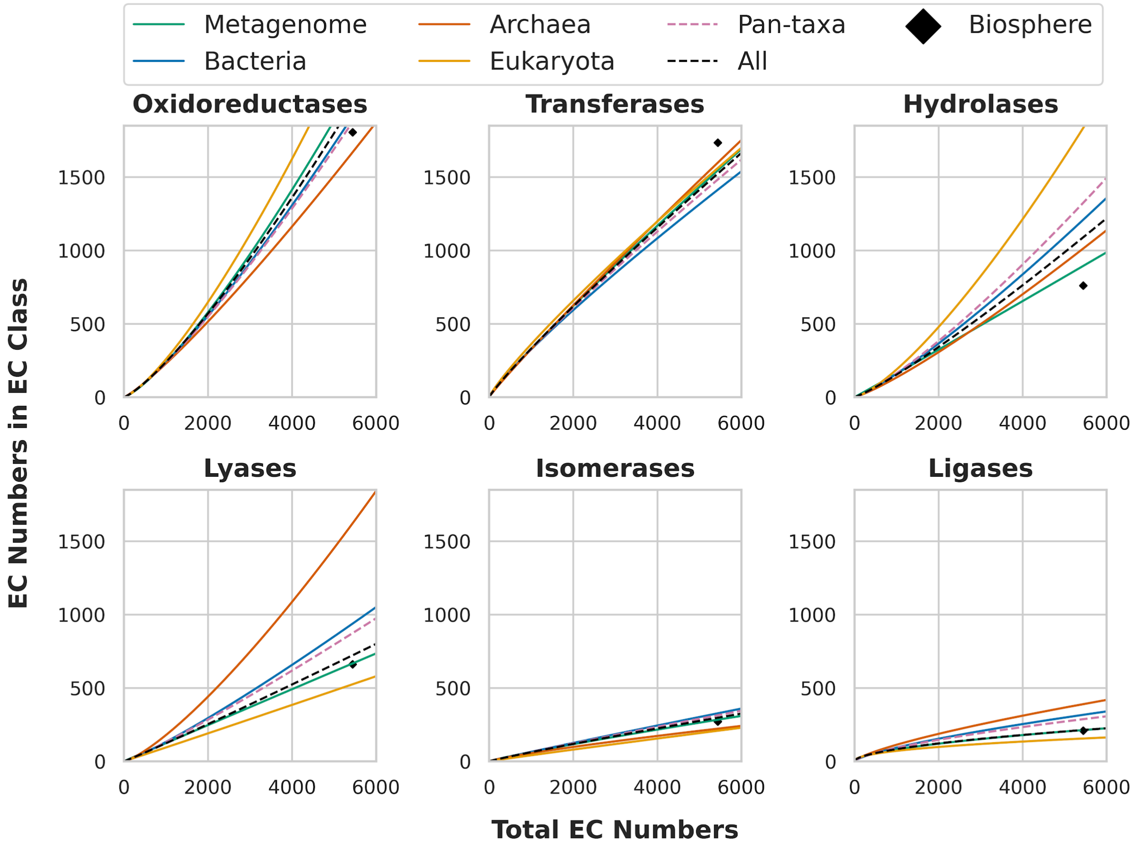


***Figure S6:*** *Scaling behaviors in enzyme functions for each EC, including scaling for data from all three domains taken together (Pan-taxa) and all domain data together also with metagneomes (All). Biosphere values are 5440 total number of EC numbers with an enzyme class breakdown of 1805 oxidoreductases, 1733 transferases, 761 hydrolases, 661 lyases, 271 isomerases, and 209 ligases.*

We also include here scaling plots with histograms showing the density of each dataset along the x and y-axes for each EC class, as shown in Figures S7 - S12.

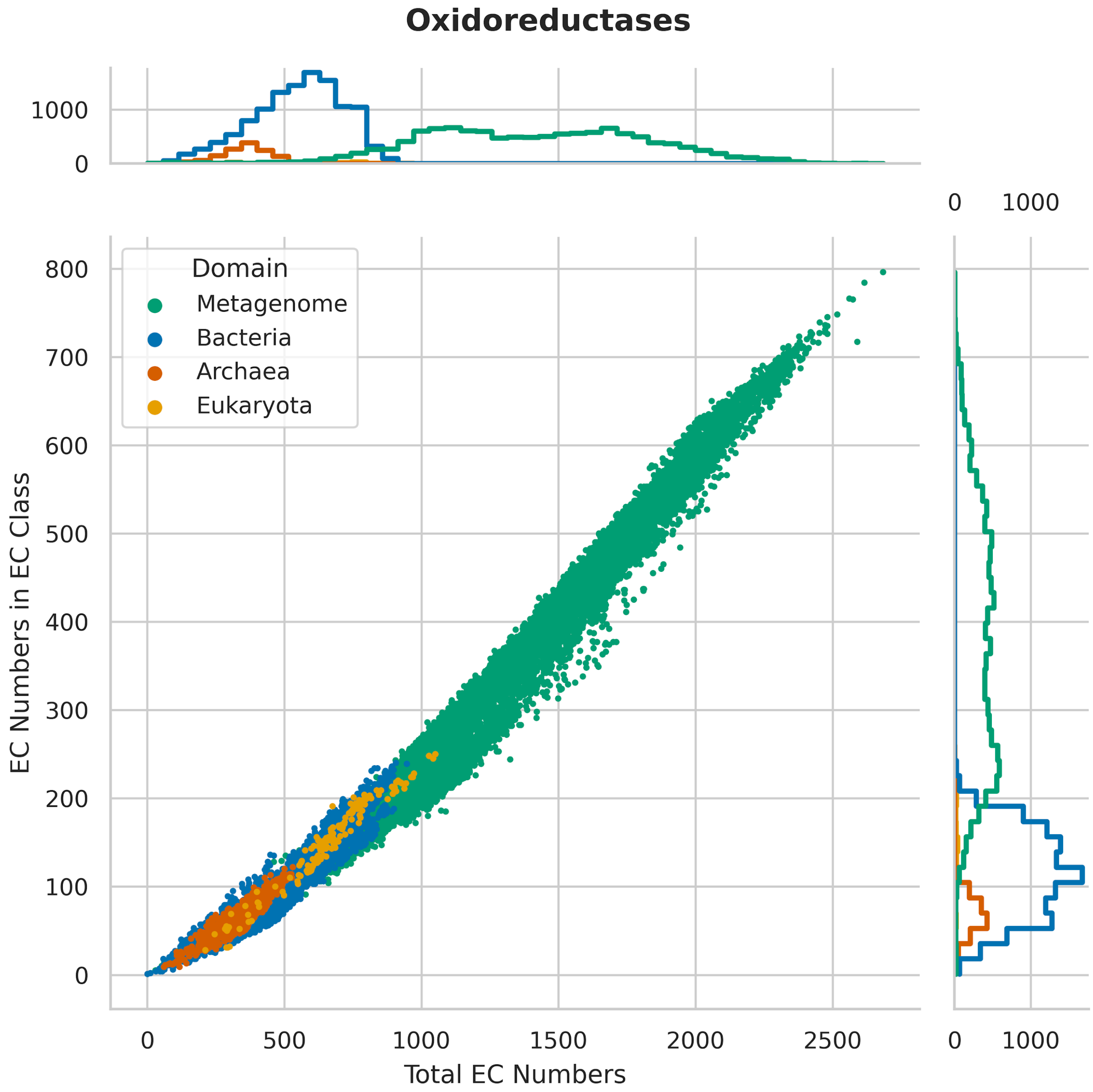


***Figure S7:*** *Distribution of Bacteria, Archaea, and Eukaryota taxa and metagenome data used for scaling plots in Figure 2 of paper for oxidoreductases. Marginal histogram plots show counts with bin width of 1.*

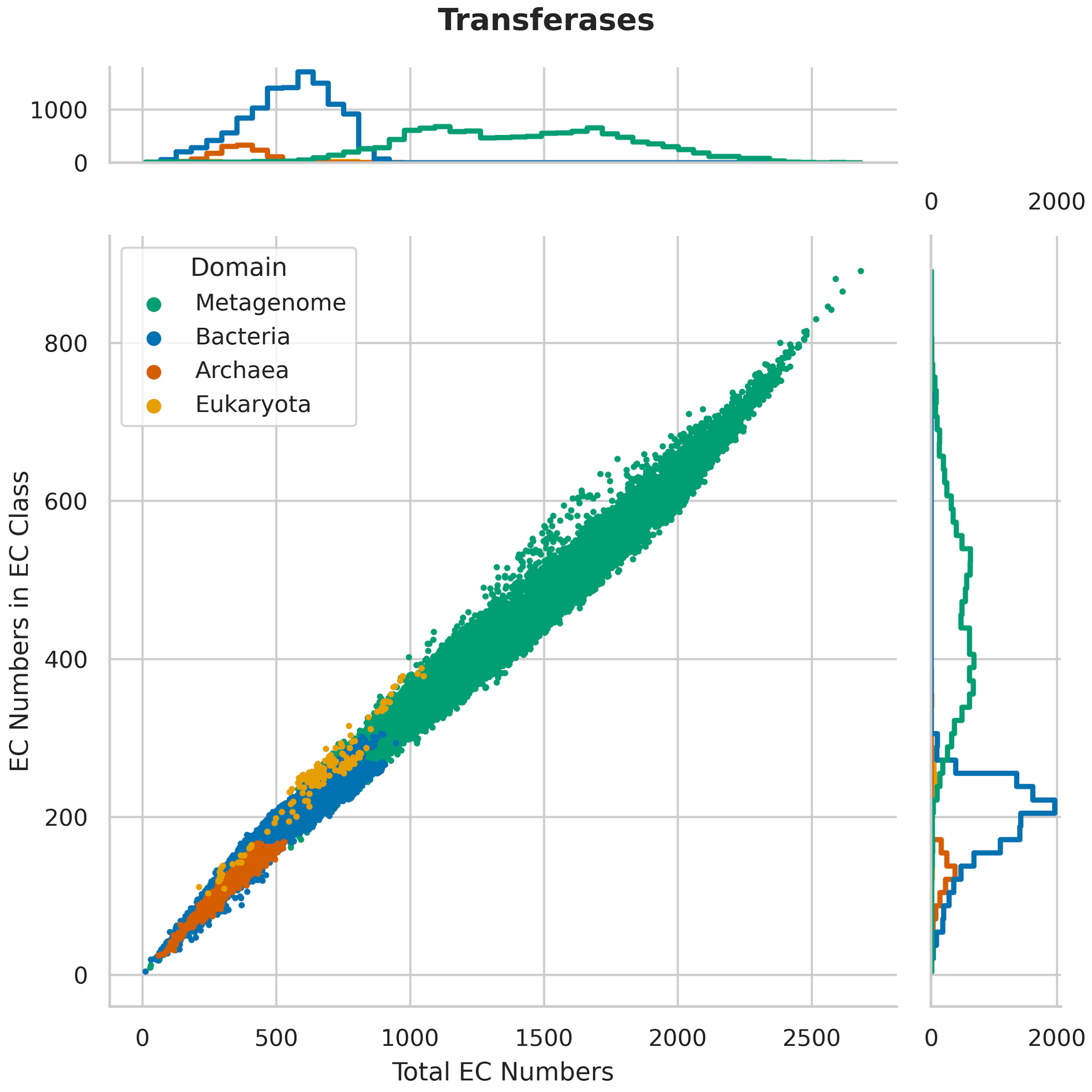


***Figure S8:*** *Distribution of Bacteria, Archaea, and Eukaryota taxa and metagenome data used for scaling plots in Figure 2 of paper for transferases. Marginal histogram plots show counts with bin width of 1.*

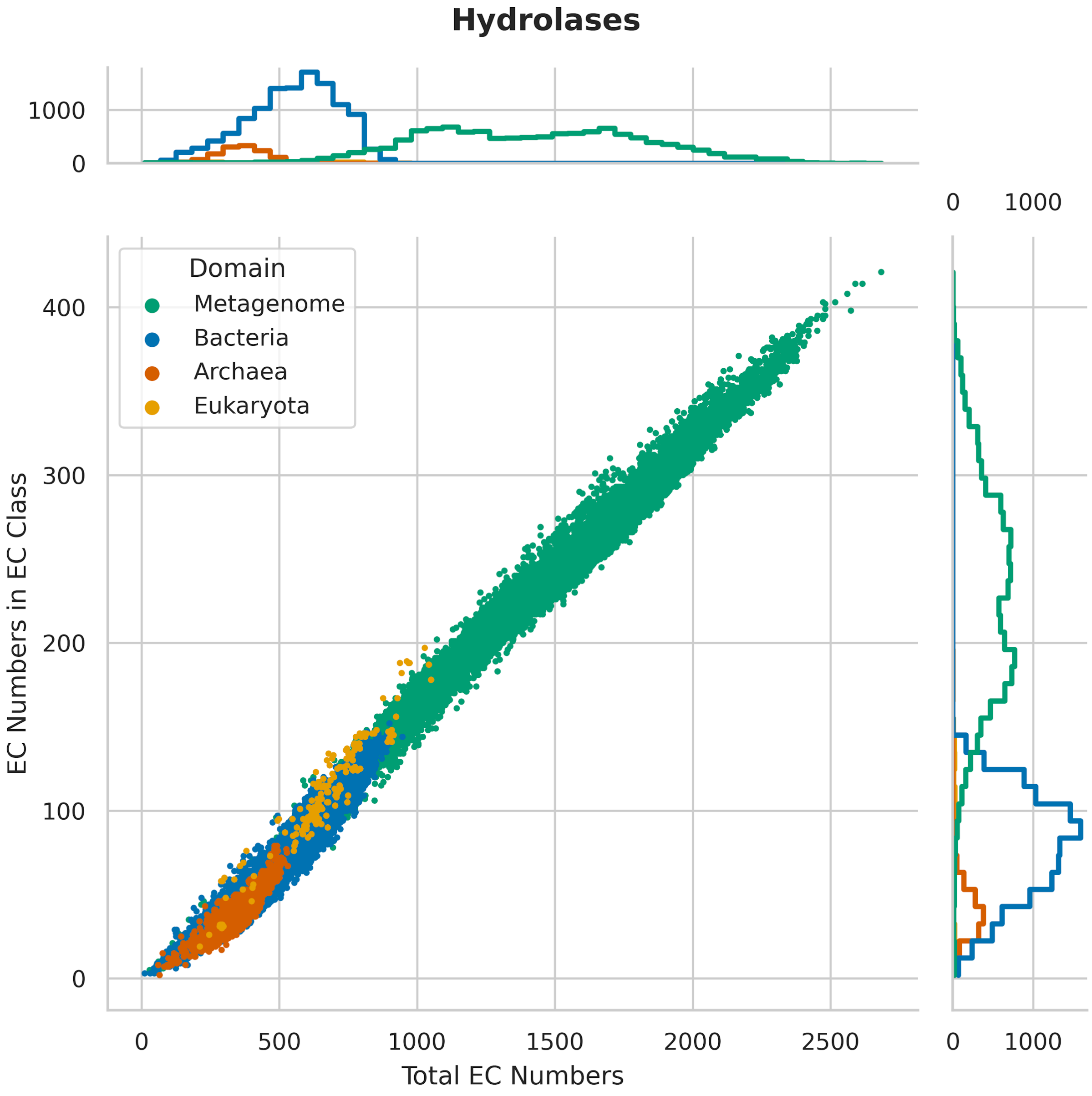


***Figure S9:*** *Distribution of Bacteria, Archaea, and Eukaryota taxa and metagenome data used for scaling plots in Figure 2 of paper for hydrolases. Marginal histogram plots show counts with bin width of 1.*

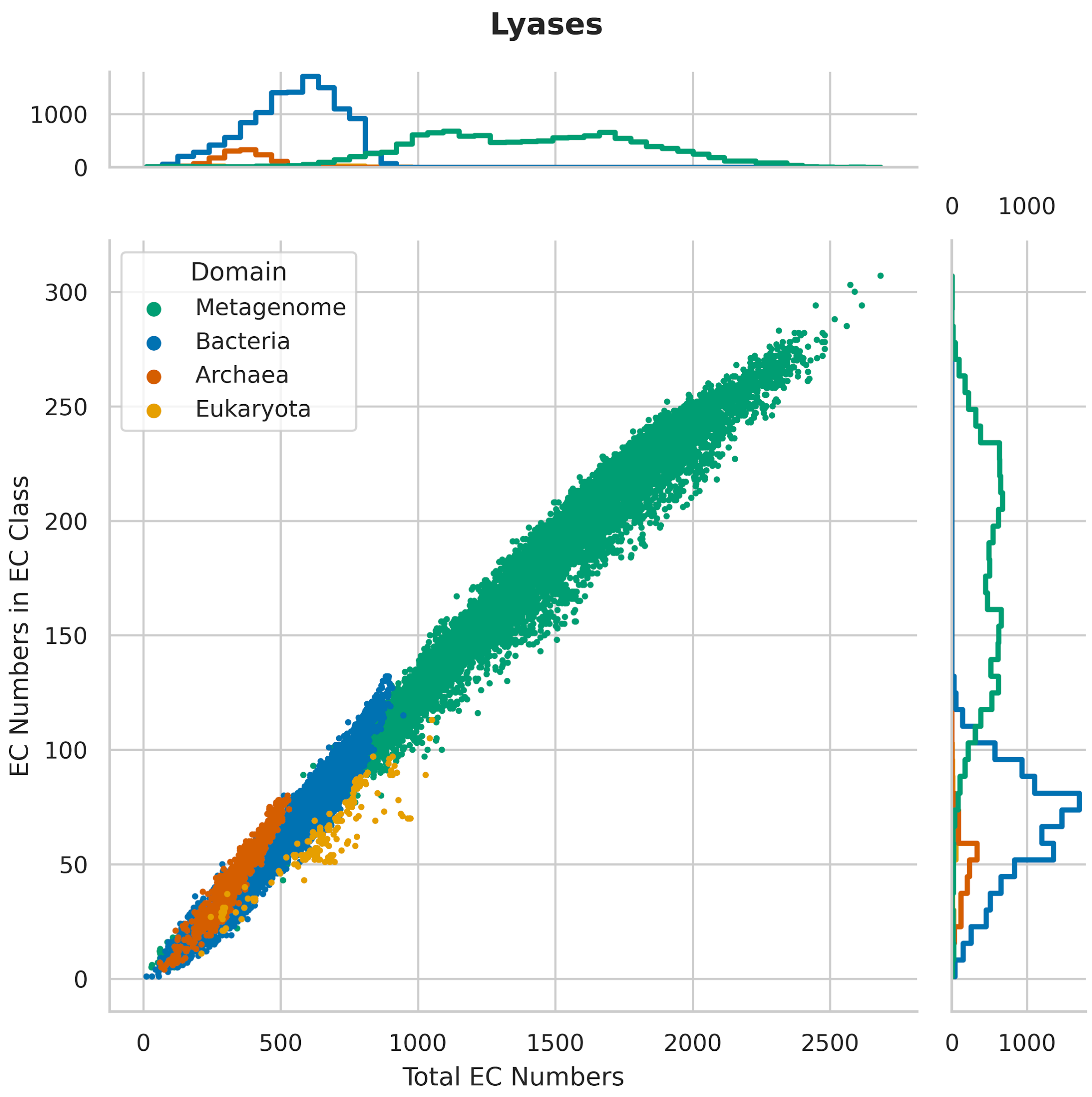


***Figure S10:*** *Distribution of Bacteria, Archaea, and Eukaryota taxa and metagenome data used for scaling plots in Figure 2 of paper for lyases. Marginal histogram plots show counts with bin width of 1.*

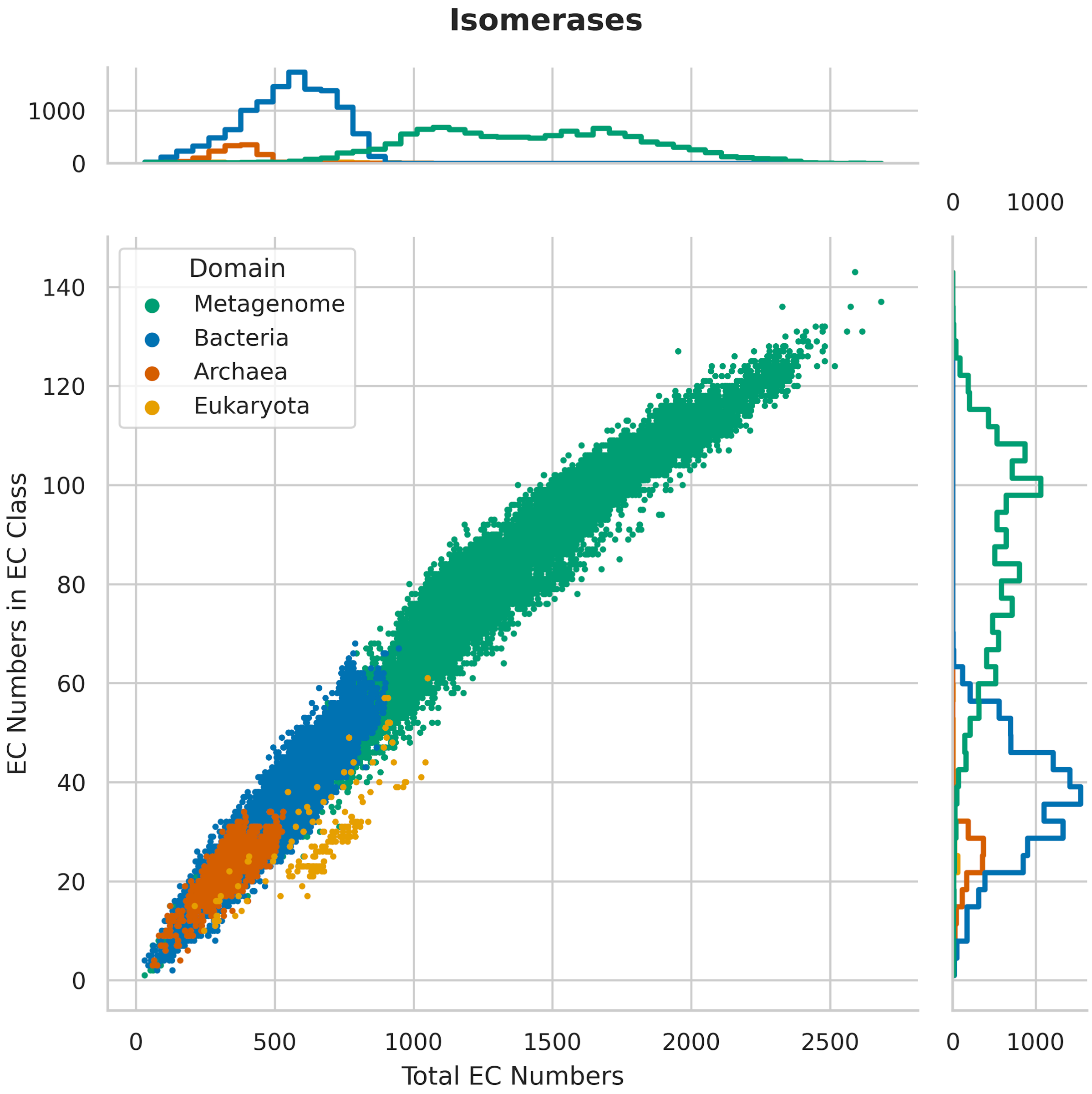


***Figure S11:*** *Distribution of Bacteria, Archaea, and Eukaryota taxa and metagenome data used for scaling plots in Figure 2 of paper for isomerases. Marginal histogram plots show counts with bin width of 1.*

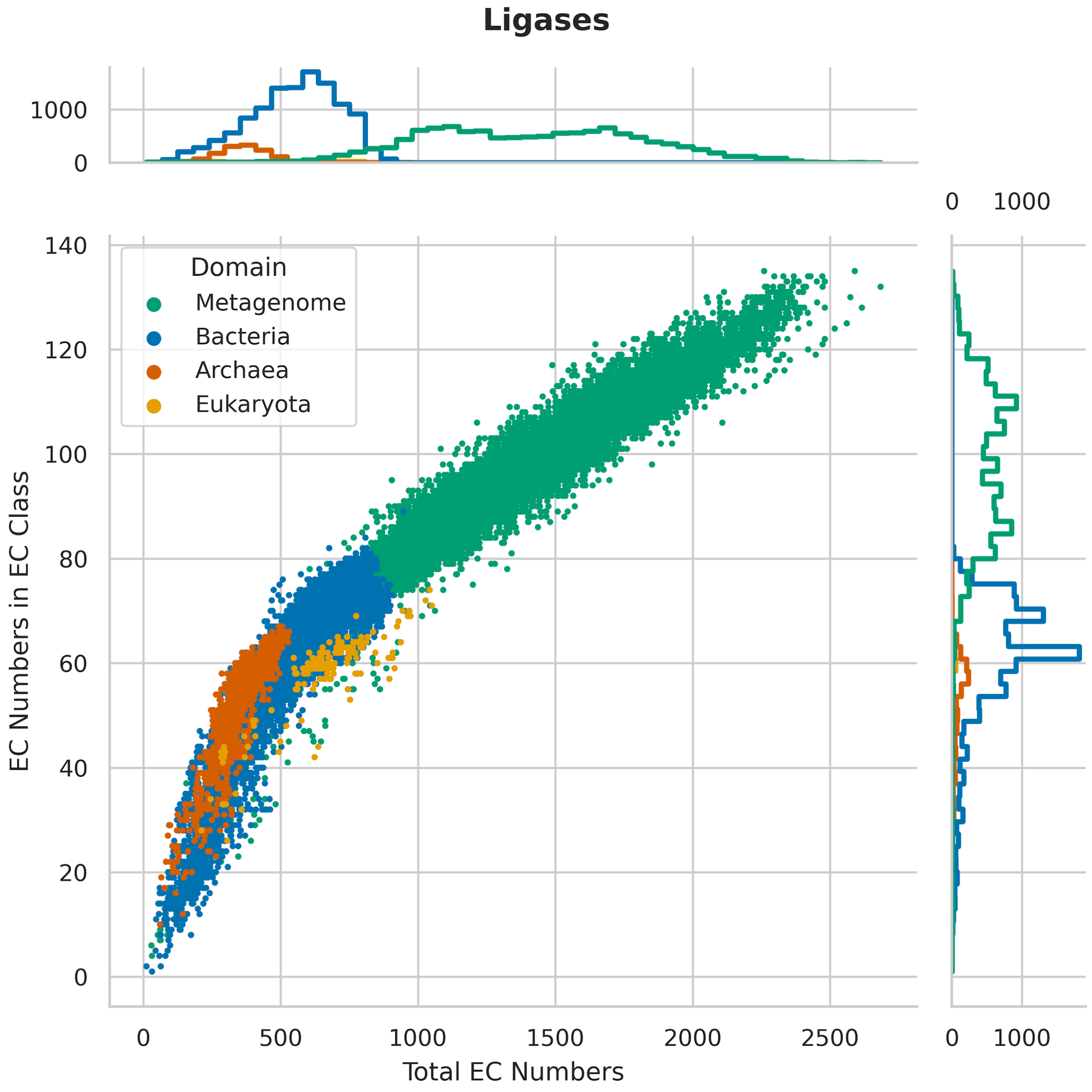


***Figure S12:*** *Distribution of Bacteria, Archaea, and Eukaryota taxa and metagenome data used for scaling plots in Figure 2 of paper for ligases. Marginal histogram plots show counts with bin width of 1.*

#### Universality of Enzyme Functions, Reactions and Compounds

To quantify universality across enzyme functions, reactions, and compounds, we calculated AUC scores for the corresponding ranked-frequency distribution curve, of each respective function, reaction or compound across each dataset. We applied Simpson's rule for the calculation of AUC score. Simpson's rule, or a three-point rule Newton-Cotes formula, is one of the techniques of numerical integration using a piecewise quadratic polynomial for approximating the area under a given arbitrary curve. We used the module scipy.integrate.simps of the Python package SciPy for the implementation of Simpson's rule.

The total number of enzyme functions, reactions and compounds in the filtered data used to calculate AUC scores are in Table S7.

| | **Total # of Unique** | | |
|---|---|---|---|
| | **Enzyme functions** | **Reactions** | **Compounds** |
| **Archaea** | 1610 | 3027 | 2826 |
| **Bacteria** | 2426 | 4259 | 3761 |
| **Eukaryota** | 2103 | 3971 | 3628 |
| **Metagenome** | 2995 | 5167 | 4536 |
| **Pan-taxa** | 2911 | 5040 | 4425 |
| **ALL** | 2998 | 5171 | 4540 |

***Table S7:*** *Table of total number of unique ECs, reactions and compounds for each dataset after filtering.*

The percentage of each EC in a given data set is shown in Figure S13 (averages and standard deviation for datasets that are ensembles of genomes or metagenomes).

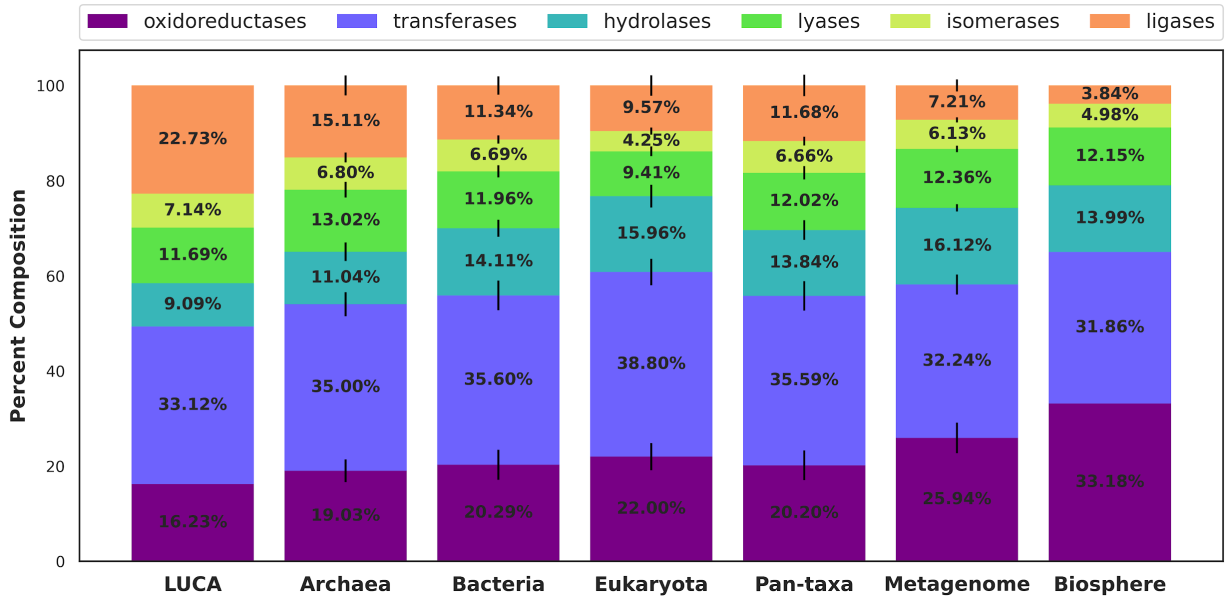


***Figure S13:*** *Percentage of ECs for the LUCA model and the Biosphere, as well as averages over ensembles for each of the data sets sampled from the three domains and metagenomes. Pan-taxa data is the ensemble average over data from the three domains. From left to right the total number of unique enzyme functions for each dataset is: 154 in the LUCA model, 1610 in archaea, 2426 in bacteria, 2103 in eukaryota, 2911 in pan-taxa, 2995 in metagenomes, and 5440 in biosphere. The data show a general trend that certain EC classes become enriched and others depleted as the total diversity of enzymes in a biochemical system increases (e.g. confirms that the scaling trends we see are held up in terms of averages for datasets).*

### Scaling Data

To determine the correlation between universality of component membership (enzyme functions, reactions, compounds) and how tightly constrained the identified EC scaling laws are, we plotted variation in the fitted value of the scaling coefficients across taxa, against the average AUC score of the three domains. Variation was calculated by taking the difference between the largest fitted value for the scaling coefficient and the smallest. The results are shown in Figure S14, which demonstrate that the distribution of enzyme functions across the data is not correlated with how tightly constrained scaling coefficients are across data sets.

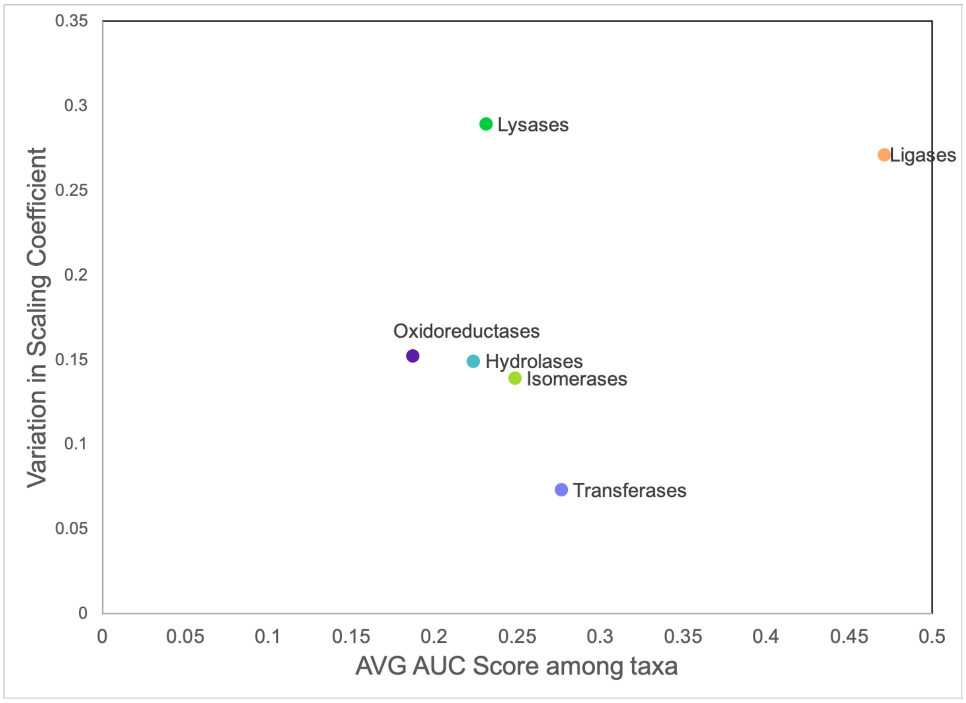


***Figure S14:*** *Variation in the scaling coefficient across taxa datasets (measuring the difference between the largest and smallest coefficient value across the domains) does not correlate with the average Area Under the Curve universality score when compared across the enzyme classes. The lack of correlation suggests that variation in scaling behavior is not directly correlated with universality of individual enzyme commission number identifiers within an Enzyme Class (EC).*

*Committee of the International Union of Biochemistry and Molecular Biology on the Nomenclature and Classification of Enzymes*. (Academic Press, 1992).

6. McDonald, A. G., Boyce, S. & Tipton, K. F. ExplorEnz: the primary source of the IUBMB enzyme list. *Nucleic Acids Res.* **37**, D593–7 (2009).
7. Hägerhäll, C. Succinate: quinone oxidoreductases. Variations on a conserved theme. *Biochim. Biophys. Acta* **1320**, 107–141 (1997).
8. Sellés Vidal, L., Kelly, C. L., Mordaka, P. M. & Heap, J. T. Review of NAD(P)H-dependent oxidoreductases: Properties, engineering and application. *Biochim. Biophys. Acta: Proteins Proteomics* **1866**, 327–347 (2018).
9. Jeske, L., Placzek, S., Schomburg, I., Chang, A. & Schomburg, D. BRENDA in 2019: a European ELIXIR core data resource. *Nucleic Acids Res.* **47**, D542–D549 (2019).
10. White, G. F., Russell, N. J. & Tidswell, E. C. Bacterial scission of ether bonds. *Microbiol. Rev.* **60**, 216–232 (1996).
11. Lipmann, F. Metabolic generation and utilization of phosphate bond energy. *Adv. Enzymol. Relat. Areas Mol. Biol.* **1**, 99–162 (1941).
12. Cuesta, S. M., Rahman, S. A. & Thornton, J. M. Exploring the chemistry and evolution of the isomerases. *Proc. Natl. Acad. Sci. U. S. A.* **113**, 1796–1801 (2016).
13. Martinez Cuesta, S., Furnham, N., Rahman, S. A., Sillitoe, I. & Thornton, J. M. The evolution of enzyme function in the isomerases. *Curr. Opin. Struct. Biol.* **26**, 121–130 (2014).
14. Hernández, S. B. & Cava, F. Environmental roles of microbial amino acid racemases. *Environ. Microbiol.* **18**, 1673–1685 (2016).
15. Holliday, G. L., Rahman, S. A., Furnham, N. & Thornton, J. M. Exploring the biological

and chemical complexity of the ligases. *J. Mol. Biol.* **426**, 2098–2111 (2014).

16. Giovannoni, S. J. *et al.* Genome streamlining in a cosmopolitan oceanic bacterium. *Science* **309**, 1242–1245 (2005).
17. Dietrich, F. S. *et al.* The Ashbya gossypii genome as a tool for mapping the ancient Saccharomyces cerevisiae genome. *Science* **304**, 304–307 (2004).
18. Harris, A. J. & Goldman, A. D. Phylogenetic reconstruction shows independent evolutionary origins of mitochondrial transcription factors from an ancient family of RNA methyltransferase proteins. *J. Mol. Evol.* **86**, 277–282 (2018).
19. Mirkin, B. G., Fenner, T. I., Galperin, M. Y. & Koonin, E. V. Algorithms for computing parsimonious evolutionary scenarios for genome evolution, the last universal common ancestor and dominance of horizontal gene transfer in the evolution of prokaryotes. *BMC Evol. Biol.* **3**, 2 (2003).
20. Delaye, L., Becerra, A. & Lazcano, A. The last common ancestor: what's in a name? *Orig. Life Evol. Biosph.* **35**, 537–554 (2005).
21. Yang, S., Doolittle, R. F. & Bourne, P. E. Phylogeny determined by protein domain content. *Proc. Natl. Acad. Sci. U. S. A.* **102**, 373–378 (2005).
22. Ranea, J. A. G., Sillero, A., Thornton, J. M. & Orengo, C. A. Protein superfamily evolution and the last universal common ancestor (LUCA). *J. Mol. Evol.* **63**, 513–525 (2006).
23. Wang, M., Yafremava, L. S., Caetano-Anollés, D., Mittenthal, J. E. & Caetano-Anollés, G. Reductive evolution of architectural repertoires in proteomes and the birth of the tripartite world. *Genome Res.* **17**, 1572–1585 (2007).
24. Srinivasan, V. & Morowitz, H. J. The canonical network of autotrophic intermediary metabolism: minimal metabolome of a reductive chemoautotroph. *Biol. Bull.* **216**, 126–130

(2009).
